# Extraordinary levels of per- and polyfluoroalkyl substances (PFAS) in vertebrate animals at a New Mexico desert oasis: multiple pathways for wildlife and human exposure

**DOI:** 10.1101/2023.10.11.561778

**Authors:** Christopher C. Witt, Chauncey R. Gadek, Jean-Luc E. Cartron, Michael J. Andersen, Mariel L. Campbell, Marialejandra Castro-Farías, Ethan F. Gyllenhaal, Andrew B. Johnson, Jason L. Malaney, Kyana N. Montoya, Andrew Patterson, Nicholas T. Vinciguerra, Jessie L. Williamson, Joseph A. Cook, Jonathan L. Dunnum

**Affiliations:** Museum of Southwestern Biology, University of New Mexico, Albuquerque, New Mexico, 87131, United States of America; Department of Biology, University of New Mexico, Albuquerque, New Mexico, 87131, United States of America; Environmental Stewardship, Los Alamos National Laboratory, Los Alamos, NM 87545, United States of America; Daniel B. Stephens & Associates, Inc., 6020 Academy Road NE, Suite 100, Albuquerque, New Mexico, 87109, United States of America; New Mexico Museum of Natural History and Science, Albuquerque, New Mexico, 87104, United States of America; Eurofins Environment Testing America, West Sacramento, California 95605, United States of America; Cornell Lab of Ornithology, Cornell University, Ithaca, NY, 14850, United States of America

**Keywords:** PFAS, Biorepositories, Birds, Mammals, Hunting, Museum collections, Holloman Air Force Base

## Abstract

Per- and polyfluoroalkyl substances (PFAS) threaten human and wildlife health, but their movement through food webs remains poorly understood. Contamination of the physical environment is widespread, but particularly concentrated at military installations. Here we measured 17 PFAS in wild, free-living mammals and migratory birds at Holloman Air Force Base (AFB), New Mexico, USA, where wastewater catchment lakes form biodiverse oases. PFAS concentrations were among the highest ever reported in animal tissues, and high levels have persisted for at least three decades. The hazardous long carbon-chain form, perfluorooctanosulfonic acid (PFOS), was most abundant, with liver concentrations averaging tens of thousands of ng/g wet weight (ww), reaching as high 97,000 ng/g (ww) in a 1994 specimen of white-footed mouse (*Peromyscus leucopus*) and 38,000 ng/g ww in a duck, the American wigeon (*Mareca americana*). Perfluorohexanesulfonic acid (PFHxS) averaged thousands of ng/g ww in the livers of birds and house mice, but one order of magnitude lower in the livers of upland desert rodent species. PFAS levels were strikingly lower at control sites, even for highly mobile migratory species. Tissue concentrations were correlated within individuals, and consistently higher in liver than in muscle or blood. Twenty of 23 vertebrate species sampled at Holloman AFB were heavily contaminated, representing multiple trophic levels and microhabitats, and implicating a range of pathways for PFAS spread: ingestion of surface water, sediments, and dust; foraging on aquatic invertebrates and plants by secondary consumers; and preying upon small vertebrates by higher level consumers, including consumption of game species by hunters. Unlike in other aquatic systems, piscivory was not an important pathway of PFAS uptake. In sum, legacy PFAS at a desert wetland have permeated the local food web across a period of decades, severely contaminating resident and migrant animals, and likely exposing humans via game meat consumption and outdoor recreation.

**Five highlights:**

- A biodiverse, wetland food web at a military base is heavily contaminated with PFAS.
- Littoral-zone mice and aquatic birds had high liver PFOS, up to 97,000 ng/g ww.
- Species and ecological variation were high among 16 PFAS detected in animal tissues.
- Game species had dangerously contaminated meat and can transport it long distances.
- Biorepositories provide key temporal and spatial sampling for contaminant studies.

**Graphical abstract:** 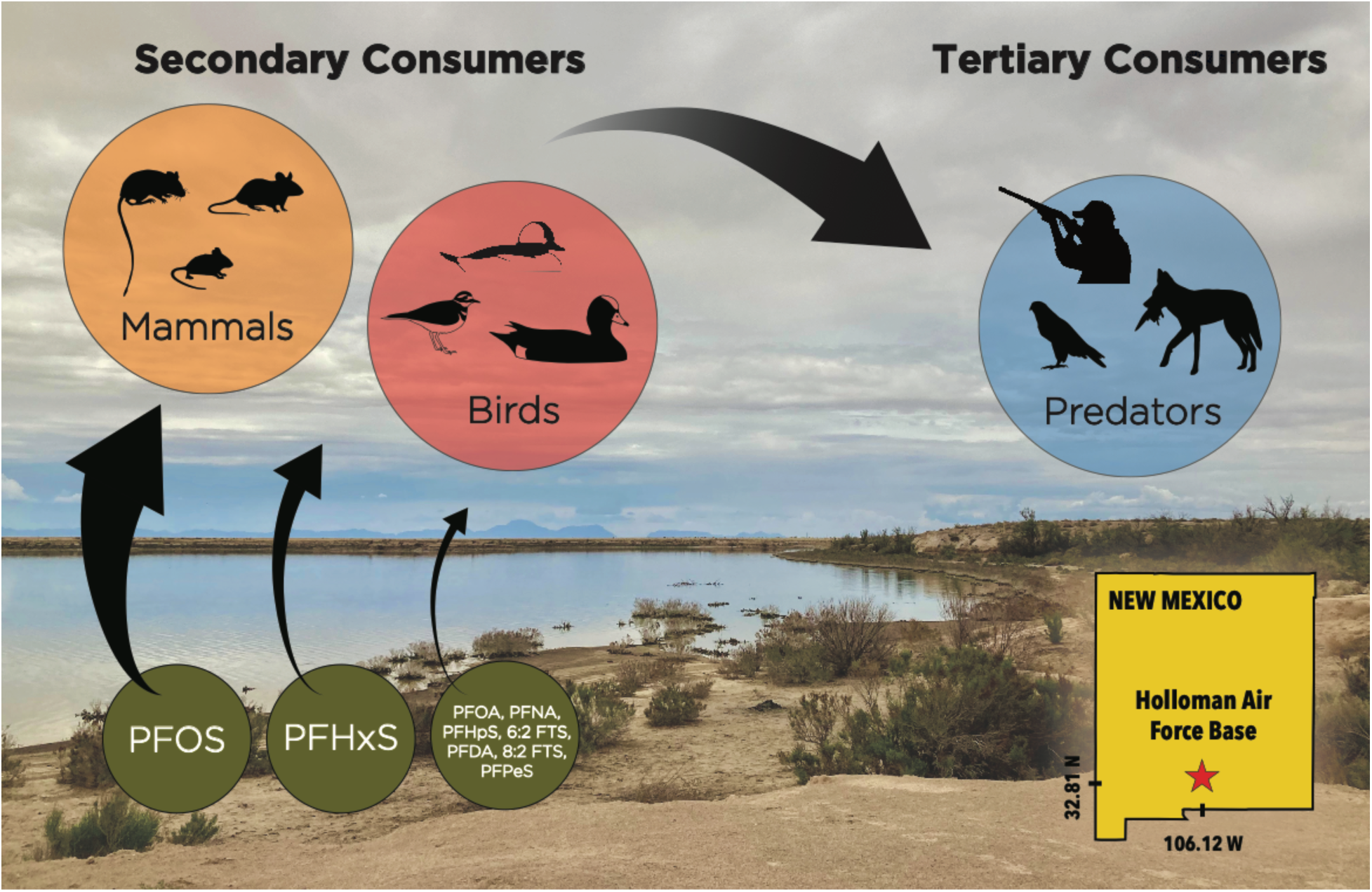

## 1. Introduction

Per- and polyfluoroalkyl substances (PFAS) are a pernicious threat to global wildlife and human health because of their long-term chemical stability, nonbiodegradability in the environment, biopersistence in tissues, and documented serious health effects (Giesy and Kannan, 2001; Lau et al., 2007; Evich et al., 2022). This group of human-made chemicals has been in widespread use since the 1940s for a variety of industrial and commercial applications, such as non-stick and stain-resistant coatings, fire-fighting foams, and waterproofing materials (US EPA, 2016; US Food and Drug Administration, 2022). Globally, military bases tend to be among the most PFAS-contaminated sites (Anderson et al., 2016). Surface and ground water on military bases are centers of accumulation and movement of PFAS substances that are known to be harmful to wildlife and human health, particularly ‘legacy’, long carbon-chain compounds that were phased out of manufacturing due to toxicity, including perfluorooctanosulfonic acid (PFOS), perfluorohexanesulfonic acid (PFHxS), and perfluorooctoanoic acid (PFOA) (Buck et al., 2011). Because of their persistence, PFAS are widespread in the environment, extending far beyond heavily polluted point sources.

Even low-level exposure has been implicated in a wide range of health and ecological impacts (Fenton et al., 2021; Grandjean and Clapp, 2015; Sebastiano et al., 2023). In the human body, modest tissue concentrations of PFAS have been linked to cancer, developmental problems, reproductive problems such as pre-term birth, autoimmune disease, and endocrine disruptions, among other serious health problems (DeWitt, 2015; Fenton et al., 2021; Taibl et al., 2023). However, movements of PFAS through food webs that include humans and domesticated and wild animals are poorly understood because of the lack of strategic and comprehensive biodiversity sampling infrastructure (Malaney and Cook, 2018) and the challenges associated with assessing exposure, sampling diverse species, and conducting precise assays (De Silva et al., 2021).

In wildlife, the evidence for negative health effects of PFAS is largely preliminary, but a growing list of studies have reported specific detrimental effects and declines in overall condition (D’Hollander et al., 2014; Costantini et al., 2019; Banyoi et al., 2022; Guillette et al., 2022; Jouanneau et al., 2022; Sebastiano et al., 2023). Toxicity estimates based on lab experiments are similar for rodents and birds; for both vertebrate classes, perfluoroalkane sulfonic acids (PFSAs) tend to be more toxic than perfluoroalkyl carboxylic acids (PFCAs) at similar carbon-chain lengths, and 8-carbon compounds tend to be more toxic than shorter molecules (Ankley et al., 2021). In birds, experimentally estimated toxicity reference values (TRV) for PFOS in blood serum and liver tissue concentrations were on the order of hundreds to thousands of parts per billion (Newsted et al., 2005); however, tissue concentrations 4–5 orders of magnitude lower in various species have been linked to endocrine disruption, immune disfunction, and low birth weights (Guillette et al., 2020; Sebastiano et al., 2023; Taibl et al., 2023).

Understanding variation in exposure, excretion, and bioaccumulation across species is a frontier for PFAS research. The toxicokinetics of various PFAS are known to be variable among species of birds and mammals and among classes of PFAS. Comparisons of PFAS levels in different tissues and among wild species at focal study sites over time can help to reveal rates and mechanisms of movement through individual organisms and food webs, as well as specific pathways of PFAS transmission that pose potential threats to animal and human health (Pizzurro et al., 2019). PFAS are proteinophilic and move easily into protein-rich tissues, such as blood and liver, where rates of bioaccumulation and trophic magnification vary profoundly among species and among classes of PFAS (Munoz et al., 2022; Ren et al., 2022; Sun et al., 2022). In general, longer carbon-chain compounds bioaccumulate faster, and PFSAs such as PFOS bioaccumulate faster than PFCAs (such as PFOA) (Conder et al., 2008).

Variable elimination rates partly underpin variable bioaccumulation rates among species. Blood serum or plasma half-lives of PFAS in varied rodent and bird species have been found to range from hours to over one year but tended to be on a scale of weeks to months, longer for PFOS than PFOA, and shorter for females that shed PFAS through egg or placental tissue (Death et al., 2021). Elimination half-lives of long carbon-chain PFAS substances tend to be much longer for humans than for other species that have been tested, ranging as high as 5.4 years for PFOS, 8.5 years for PFOA, and 15.5 years for PFHxS (Pizzurro et al., 2019). An example of species-specific biokinetics is provided by a recent comparative study between co-occurring great tits *(Parus major)* and blue tits *(Cyanistes caeruleus)* across a gradient of exposure, showing that even closely related species can differ in how they sequester long carbon-chain versus short carbon-chain perfluoroalkyl carboxylic acids (PFCAs) (Lasters et al., 2021). At higher-level taxonomic scales, terrestrial plants and herbivorous invertebrates tended to sequester more short-chain PFCAs, whereas vertebrate animals and higher trophic level invertebrates tended to sequester more long-chain PFCAs (Groffen et al., 2022).

Holloman Air Force Base and its immediate vicinity, in the Chihuahuan Desert of southcentral New Mexico, contain artificial wetlands contaminated with PFOS, PFHxS, PFOA, and other PFAS (Jarvis et al., 2021). Holloman Lake was created in the 1965 by installing an earthen dam on a playa lake (ephemeral wetland). The lake receives stormwater runoff and treated sewage from Holloman Air Force Base. Many contaminants accumulated over decades of wastewater deposition, among which were components of aqueous film-forming foams (AFFF) that had been used extensively in fire-fighting training since the 1970s (Moody and Field, 2000). While some components of the various AFFF formulations degrade to PFOA and other perfluoroalkyl carboxylic acids (PFCAs), others degrade to PFOS and other perfluoroalkyl sulfonic acids (PFSAs); however, the ingredients of proprietary AFFF formulations and their pathways of degradation in the environment are not completely known (Anderson et al., 2016). A recent analysis of all publicly available data for surface waters across the United States showed that Holloman Lake is one of the most polluted with PFOS, with measured concentrations as high as 5.9 ng/mL and a median concentration of 4.5 ng/mL; other wetlands on and around Holloman Airforce base are also heavily contaminated (Jarvis et al., 2021).

The wetlands, riparian, and littoral habitat surrounding Holloman Lake are largely managed by the U. S. Department of Defense, the U. S. Department of Interior (Bureau of Land Management), and the New Mexico State Lands Office. This biodiverse ecosystem includes a woodland area at the head of the lake, tall emergent vegetation, mudflats, permanent open water, muddy shorelines, and extensive desert shrubland surrounding the lake. Holloman Lake the largest and most ecologically significant water source in the entire Tularosa Basin (∼16,800 km^2^). The dominant plants of the lake’s shoreline are small trees (<4 m) and shrubs, the vast majority of which are tamarisk (*Tamarix* sp.) and four-wing saltbush (*Atriplex canescens*). Western glasswort (*Salicornia rubra*), saltgrass (*Distichlis spicata*), and harmel (*Peganum* sp.) are also present. The inflow to the lake is marked by taller (<6 m) tamarisk trees and emergent aquatic vegetation such as cattail (*Typhus latifolia*) and clubrush (*Bolboschoenus* sp.), providing habitat for numerous vertebrate animal species to shelter, forage, and nest. As a result of Holloman Lake’s aquatic resources, surrounding vegetation, and inflow of nutrient rich treated sewage, the site is home to a diverse vertebrate fauna, including birds, mammals, reptiles, and amphibians. Citizen science data from eBird.org shows that at least 252 species of birds occur at Holloman Lake, 113 of which are aquatic species, and 41 of which are game species that can be legally hunted. The mammal community is also diverse and typical of the northern Chihuahuan Desert scrub community, with more than 35 non-volant mammal species occurring locally (Frey and Yates, 1996; Malaney et al., 2022) and at least 19 species of bats occurring within 100 km, some of which are long-distance migrant species (Cryan, 2003; Russell et al., 2005). Free-ranging beef cattle (Bos *taurus*) and oryx (*Oryx gazella*) graze in the vicinity and drink directly from the lake, the latter species having been introduced in the late 1960s and 1970s for sport hunting.

Hunting for sport or subsistence provides a pathway for PFAS movement from contaminated wildlife into human tissues (Haug et al., 2010), although concerns about PFAS-contaminated fish consumption have received substantially more attention (Guillette et al., 2020). In the few cases worldwide where assays have been conducted on game species, such as wild boars and wild ducks, their tissues have been shown to harbor potentially dangerous concentrations of PFAS (typically PFOS, PHFxS, PFOA, et al.) (Death et al., 2021; Rupp et al., 2023).

Only a few previous studies have published data on PFAS in waterfowl species that are subject to hunting. At Tokyo Bay in Japan, mallards (*Anas platyrhynchos*) and northern pintails (*Anas acuta*) had PFOS in liver tissue as high as ∼500 ng/g wet weight (ww) (Taniyasu et al., 2003). In Hudson Bay, Canada, apparently far from any point source, PFOS in liver tissue was in the single-digit ng/g range for common eiders (*Somateria mollissima*) and in the tens of ng/g ww for white-winged scoters (*Melanitta deglandi*), with one individual measuring up to 120 ng/g ww (Kelly et al., 2009). In southeastern Australia, four species of ducks were surveyed across 19 wetlands of which a subset had sediments and water that were contaminated with PFAS; grey teal (*Anas gracilis*) and chestnut teal (*Anas castanea*) liver tissues were contaminated with PFOS and PHFxS up to the tens of ng/g ww, while pink-eared duck (*Malacorhynchus membranaceus*) and Pacific black duck (*Anas superciliosa*) livers were found to contain hundreds of ng/g ww (maximum 340 ng/g ww; median 9.5 ng/g ww), high enough to warrant government investigations and to trigger warnings about consumption (Senversa, 2018; Environmental Protection Authority Victoria, 2019; Sharp et al., 2021). Breast muscle tissue concentrations in the same Australian duck individuals were one order of magnitude lower than in liver. Among the 19 wetlands studied, some of the most contaminated ducks were found at non-contaminated sites, suggesting dispersal after exposure, leading to uncertain implications for hunters and other upper trophic predators (Sharp et al., 2021). As in other systems, species-specific foraging habits were thought to have caused different levels of exposure to PFAS from sediment, fish, invertebrates, water, and plants (Larson et al., 2018).

In this study, we explored the extent and pathways of PFAS contamination in wild animal populations in the area of Holloman AFB (Fig. 1). Specifically, we asked (1) Do vertebrate secondary consumers at a PFAS-contaminated wetland accumulate PFAS in their tissues, providing pathways for PFAS to permeate biodiverse food webs? (2) Have PFAS been present since at least 1994, when the first museum-archived tissues samples from the area were collected? (3) Do migratory game bird species at a contaminated point source show elevated PFAS in tissues relative to control samples from uncontaminated sites, and does their harvested meat pose a potential danger to human health? (4) To what extent do terrestrial small mammal species that have no direct contact with lake water or sediments harbor PFAS in their tissues, thereby suggesting groundwater-, dust-, or aerosol-mediated pathways that could also expose humans? (5) How do PFAS profiles and tissue distributions vary across species with varying phylogenetic affinities, habitats, and diets and what does this indicate about pathways for PFAS uptake, bioaccumulation, and trophic magnification?

**Fig. 1.**
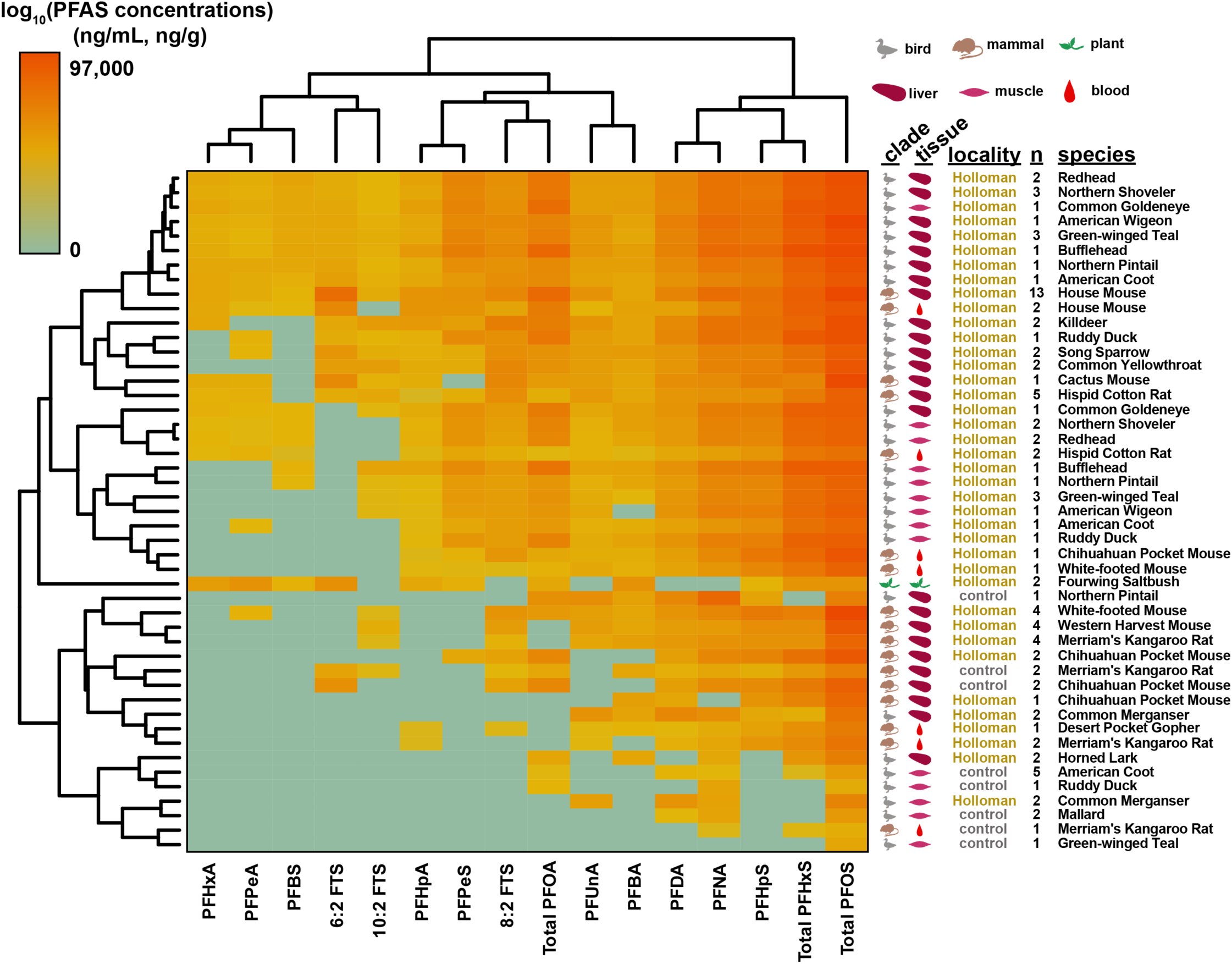
Heat map showing similarity of PFAS concentration-profiles in 47 unique species-tissue combinations from Holloman Air Force base (yellow font) versus control sites (blue font). Tissues include bird muscle (ng/g), bird and small mammal liver (ng/g), small mammal blood (ng/mL), and plant leaf and stem. Sample sizes for each species-tissue combination are indicated under the column labeled ‘n’. Dendrograms depict hierarchical relationships among species-tissue combinations (left) based on similarity of their 16 PFAS concentrations, and among the 16 PFAS (top) based on similarity of their concentrations across species-tissue categories. Samples clustered in accordance with ecology and phylogeny, as described in Results.

## 2. Materials and methods

### 2.1. Sampling and museum archiving

We screened 99 samples (n = 63 liver, 24 muscle, 10 blood, 2 leaves and stems), representing 34 individual birds, 40 mammals, and 2 composite plant samples (Table 1). The bird and mammal species were selected in part because of their varied phylogenetic affinities, foraging ecology, and habitat preferences. The sampled bird community included game and non-game bird species, aquatic and terrestrial species, as well as winter and breeding season residents; in total a representative set of 11 aquatic game bird species, 3 songbird species, and 1 shorebird species, all of which occur in or at the margins of contaminated wetlands at Holloman AFB. The 11 aquatic game species that we screened included 10 duck species (Anseriformes: Anatidae) and the American coot (Gruiformes: Rallidae), all of which vary in foraging depths, and, with the possible exception of the piscivorous common merganser (*Mergus merganser*), all of which are commonly hunted and eaten. The mammals comprised nine rodent species that are resident throughout their lives in the immediate littoral zone and/or surrounding desert habitats.

**Table 1.** Specimens screened for PFAS substances in this study, with links to their online data and metadata in the Arctos Collection Management System (arctosdb.org). Specimens are sorted by class of animal or plant, then alphabetically by English name of the species. Four mammal samples with localities marked with * were sampled at Holloman AFB in 1994, but at different specific sites than the samples that were collected in 2021–2023. Plant sample identifiers have prefix “320- “ and, unlike the animal identifiers, are not NK numbers.

| Class | Tissue | Identifier |  | Scientific name | locality | URL (link) with MSB GUID (catalog number) |
| --- | --- | --- | --- | --- | --- | --- |
|  | types | (NK) | English name |  |  |  |
| bird | muscle | 283613 | American Coot | *Fulica americana* | Control site | https://arctos.database.museum/guid/MSB:Bird:51000 |
| bird | muscle | 283614 | American Coot | *Fulica americana* | Control site | https://arctos.database.museum/guid/MSB:Bird:51001 |
| bird | muscle | 283616 | American Coot | *Fulica americana* | Control site | https://arctos.database.museum/guid/MSB:Bird:51003 |
| bird | muscle | 283676 | American Coot | *Fulica americana* | Control site | https://arctos.database.museum/guid/MSB:Bird:51040 |
| bird | muscle | 283697 | American Coot | *Fulica americana* | Control site | https://arctos.database.museum/guid/MSB:Bird:51044 |
| bird | liver, muscle | 283603 | American Coot | *Fulica americana* | Holloman AFB | https://arctos.database.museum/guid/MSB:Bird:50990 |
| bird | liver, muscle | 283623 | American Wigeon | *Mareca americana* | Holloman AFB | https://arctos.database.museum/guid/MSB:Bird:51010 |
| bird | liver, muscle | 283693 | Bufflehead | *Bucephala albeola* | Holloman AFB | https://arctos.database.museum/guid/MSB:Bird:51169 |
| bird | liver, muscle | 283609 | Common Goldeneye | *Bucephala clangula* | Holloman AFB | https://arctos.database.museum/guid/MSB:Bird:50996 |
| bird | liver, muscle | 283612 | Common Merganser | *Mergus merganser* | Holloman AFB | https://arctos.database.museum/guid/MSB:Bird:50999 |
| bird | liver, muscle | 283637 | Common Merganser | *Mergus merganser* | Holloman AFB | https://arctos.database.museum/guid/MSB:Bird:51024 |
| bird | liver | 283754 | Common Yellowthroat | *Geothlypis trichas* | Holloman AFB | https://arctos.database.museum/guid/MSB:Bird:51156 |
| bird | liver | 283756 | Common Yellowthroat | *Geothlypis trichas* | Holloman AFB | https://arctos.database.museum/guid/MSB:Bird:51157 |
| bird | muscle | 283617 | Green-winged Teal | *Anas crecca* | Control site | https://arctos.database.museum/guid/MSB:Bird:51004 |
| bird | liver, muscle | 283634 | Green-winged Teal | *Anas crecca* | Holloman AFB | https://arctos.database.museum/guid/MSB:Bird:51021 |
| bird | liver, muscle | 283635 | Green-winged Teal | *Anas crecca* | Holloman AFB | https://arctos.database.museum/guid/MSB:Bird:51022 |
| bird | liver, muscle | 283675 | Green-winged Teal | *Anas crecca* | Holloman AFB | https://arctos.database.museum/guid/MSB:Bird:51039 |
| bird | liver | 283645 | Horned Lark | *Eremophila alpestris* | Holloman AFB | https://arctos.database.museum/guid/MSB:Bird:50946 |
| bird | liver | 283648 | Horned Lark | *Eremophila alpestris* | Holloman AFB | https://arctos.database.museum/guid/MSB:Bird:50972 |
| bird | liver | 283771 | Killdeer | *Charadrius vociferus* | Holloman AFB | https://arctos.database.museum/guid/MSB:Bird:51309 |
| bird | liver | 283770 | Killdeer | *Charadrius vociferus* | Holloman AFB | https://arctos.database.museum/guid/MSB:Bird:51310 |
| bird | muscle | 283706 | Mallard | *Anas platyrhynchos* | Control site | https://arctos.database.museum/guid/MSB:Bird:51046 |
| bird | muscle | 283721 | Mallard | *Anas platyrhynchos* | Control site | https://arctos.database.museum/guid/MSB:Bird:51121 |
| bird | liver, muscle | 283604 | Northern Pintail | *Anas acuta* | Holloman AFB | https://arctos.database.museum/guid/MSB:Bird:50991 |
| bird | liver | 283752 | Northern Pintail | *Anas acuta* | Control site | https://arctos.database.museum/guid/MSB:Bird:51118 |
| bird | liver, muscle | 283610 | Northern Shoveler | *Spatula clypeata* | Holloman AFB | https://arctos.database.museum/guid/MSB:Bird:50997 |
| bird | liver, muscle | 283666 | Northern Shoveler | *Spatula clypeata* | Holloman AFB | https://arctos.database.museum/guid/MSB:Bird:51032 |
| bird | liver | 284405 | Northern Shoveler | *Spatula clypeata* | Holloman AFB | https://arctos.database.museum/guid/MSB:Bird:51518 |
| bird | liver, muscle | 283628 | Redhead | *Aythya americana* | Holloman AFB | https://arctos.database.museum/guid/MSB:Bird:51015 |
| bird | liver, muscle | 283668 | Redhead | *Aythya americana* | Holloman AFB | https://arctos.database.museum/guid/MSB:Bird:51034 |
| bird | muscle | 283699 | Ruddy Duck | *Oxyura jamaicensis* | Control site | https://arctos.database.museum/guid/MSB:Bird:51167 |
| bird | liver, muscle | 283630 | Ruddy Duck | *Oxyura jamaicensis* | Holloman AFB | https://arctos.database.museum/guid/MSB:Bird:51017 |
| bird | liver | 283680 | Song Sparrow | *Melospiza melodia* | Holloman AFB | https://arctos.database.museum/guid/MSB:Bird:51062 |
| bird | liver | 284404 | Song Sparrow | *Melospiza melodia* | Holloman AFB | https://arctos.database.museum/guid/MSB:Bird:51519 |
| mammal | liver | 311395 | Cactus mouse | *Peromyscus eremicus* | Holloman AFB | https://arctos.database.museum/guid/MSB:Mamm:340090 |
| mammal | liver | 311437 | Chihuahuan pocket mouse | *Chaetodipus eremicus* | Control site | https://arctos.database.museum/guid/MSB:Mamm:340081 |
| mammal | liver | 311435 | Chihuahuan pocket mouse | *Chaetodipus eremicus* | Control site | https://arctos.database.museum/guid/MSB:Mamm:340107 |
| mammal | liver | 310892 | Chihuahuan pocket mouse | *Chaetodipus eremicus* | Holloman AFB | https://arctos.database.museum/guid/MSB:Mamm:341583 |
| mammal | liver | 310831 | Chihuahuan pocket mouse | *Chaetodipus eremicus* | Holloman AFB | https://arctos.database.museum/guid/MSB:Mamm:341589 |
| mammal | blood | 311411 | Desert pocket gopher | *Geomys arenarius* | Holloman AFB | https://arctos.database.museum/guid/MSB:Mamm:340117 |
| mammal | liver | 311397 | Hispid cotton rat | *Sigmodon hispidus* | Holloman AFB | https://arctos.database.museum/guid/MSB:Mamm:340128 |
| mammal | liver | 310873 | Hispid cotton rat | *Sigmodon hispidus* | Holloman AFB | https://arctos.database.museum/guid/MSB:Mamm:341543 |
| mammal | liver, blood | 310903 | Hispid cotton rat | *Sigmodon hispidus* | Holloman AFB | https://arctos.database.museum/guid/MSB:Mamm:342181 |
| mammal | liver, blood | 310945 | Hispid cotton rat | *Sigmodon hispidus* | Holloman AFB | https://arctos.database.museum/guid/MSB:Mamm:342224 |
| mammal | liver | 31808 | Hispid cotton rat | *Sigmodon hispidus* | Holloman AFB* | https://arctos.database.museum/guid/MSB:Mamm:91660 |
| mammal | liver | 311426 | House mouse | *Mus musculus* | Holloman AFB | https://arctos.database.museum/guid/MSB:Mamm:340078 |
| mammal | liver | 311423 | House mouse | *Mus musculus* | Holloman AFB | https://arctos.database.museum/guid/MSB:Mamm:340102 |
| mammal | liver, blood | 311424 | House mouse | *Mus musculus* | Holloman AFB | https://arctos.database.museum/guid/MSB:Mamm:340103 |
| mammal | liver | 311422 | House mouse | *Mus musculus* | Holloman AFB | https://arctos.database.museum/guid/MSB:Mamm:340121 |
| mammal | liver | 310884 | House mouse | *Mus musculus* | Holloman AFB | https://arctos.database.museum/guid/MSB:Mamm:341567 |
| mammal | liver | 310883 | House mouse | *Mus musculus* | Holloman AFB | https://arctos.database.museum/guid/MSB:Mamm:341605 |
| mammal | liver | 310882 | House mouse | *Mus musculus* | Holloman AFB | https://arctos.database.museum/guid/MSB:Mamm:341606 |
| mammal | liver | 311886 | House mouse | *Mus musculus* | Holloman AFB | https://arctos.database.museum/guid/MSB:Mamm:342156 |
| mammal | liver | 311887 | House mouse | *Mus musculus* | Holloman AFB | https://arctos.database.museum/guid/MSB:Mamm:342170 |
| mammal | liver | 310922 | House mouse | *Mus musculus* | Holloman AFB | https://arctos.database.museum/guid/MSB:Mamm:342193 |
| mammal | liver | 310912 | House mouse | *Mus musculus* | Holloman AFB | https://arctos.database.museum/guid/MSB:Mamm:342196 |
| mammal | liver, blood | 310921 | House mouse | *Mus musculus* | Holloman AFB | https://arctos.database.museum/guid/MSB:Mamm:342209 |
| mammal | liver | 310959 | House mouse | *Mus musculus* | Holloman AFB | https://arctos.database.museum/guid/MSB:Mamm:342214 |
| mammal | blood | 310855 | Chih. grasshopper mouse | *Onychomys arenicola* | Holloman AFB | https://arctos.database.museum/guid/MSB:Mamm:341485 |
| mammal | liver | 310904 | Chih. grasshopper mouse | *Onychomys arenicola* | Holloman AFB | https://arctos.database.museum/guid/MSB:Mamm:342188 |
| mammal | liver | 311406 | Merriam's kangaroo rat | *Dipodomys merriami* | Holloman AFB | https://arctos.database.museum/guid/MSB:Mamm:340136 |
| mammal | liver | 310840 | Merriam's kangaroo rat | *Dipodomys merriami* | Holloman AFB | https://arctos.database.museum/guid/MSB:Mamm:341596 |
| mammal | liver, blood | 310911 | Merriam's kangaroo rat | *Dipodomys merriami* | Holloman AFB | https://arctos.database.museum/guid/MSB:Mamm:342148 |
| mammal | liver, blood | 310910 | Merriam's kangaroo rat | *Dipodomys merriami* | Holloman AFB | https://arctos.database.museum/guid/MSB:Mamm:342187 |
| mammal | liver, blood | 310941 | Merriam's kangaroo rat | *Dipodomys merriami* | Control site | https://arctos.database.museum/guid/MSB:Mamm:342206 |
| mammal | liver | 310939 | Merriam's kangaroo rat | *Dipodomys merriami* | Control site | https://arctos.database.museum/guid/MSB:Mamm:342231 |
| mammal | liver | 311390 | Western Harvest Mouse | *Reithrodontomys megalotis* | Holloman AFB | https://arctos.database.museum/guid/MSB:Mamm:340087 |
| mammal | liver | 311891 | Western Harvest Mouse | *Reithrodontomys megalotis* | Holloman AFB | https://arctos.database.museum/guid/MSB:Mamm:342158 |
| mammal | liver | 310963 | Western Harvest Mouse | *Reithrodontomys megalotis* | Holloman AFB | https://arctos.database.museum/guid/MSB:Mamm:342221 |
| mammal | liver | 31807 | Western Harvest Mouse | *Reithrodontomys megalotis* | Holloman AFB* | https://arctos.database.museum/guid/MSB:Mamm:91659 |
| mammal | liver | 310837 | White-footed mouse | *Peromyscus leucopus* | Holloman AFB | https://arctos.database.museum/guid/MSB:Mamm:341472 |
| mammal | liver, blood | 310914 | White-footed mouse | *Peromyscus leucopus* | Holloman AFB | https://arctos.database.museum/guid/MSB:Mamm:342191 |
| mammal | liver | 10440 | White-footed mouse | *Peromyscus leucopus* | Holloman AFB* | https://arctos.database.museum/guid/MSB:Mamm:89197 |
| mammal | liver | 31806 | White-footed mouse | *Peromyscus leucopus* | Holloman AFB* | https://arctos.database.museum/guid/MSB:Mamm:92667 |
| dicot | mixed | 100716-1 | Four-wing saltbush | *Atriplex canescens* | Holloman AFB | N/A |
| dicot | mixed | 100716-2 | Four-wing saltbush | *Atriplex canescens* | Holloman AFB | N/A |

Four mammal specimens were collected in the vicinity of a golf course on Holloman AFB, near a contaminated wetland, Lagoon G, at Holloman AFB, Otero County, New Mexico in 1994. Livers from these specimens were cryogenically preserved in liquid nitrogen and archived at the Museum of Southwestern Biology (MSB), University of New Mexico. We were not granted access to resample sites on Holloman AFB.

The majority of samples in this study were field collected during 2021–2023 adjacent to Holloman AFB, with most of the samples taken near Holloman Lake (32.81 N, 106.12 W ± 2 km). All historical and contemporary samples Holloman AFB were collected within a ∼2 km radius. To provide comparative ‘control’ samples, additional specimens were collected from uncontaminated sites ∼10–150 km from Holloman AFB (see Table 1).

Four small mammal specimens were collected on Holloman Air Force Base in 1994, with frozen liver samples stored in the Museum of Southwestern Biology. We screened these four samples, collected within 2 km of our recent sampling, in order to gain insights into historical contamination of wildlife in the area. These four samples were collected in the vicinity of a golf course on Holloman AFB, near a contaminated wetland, Lagoon G, to which we were not granted access for resampling.

Birds were collected by licensed hunters using shotguns with non-toxic shot, operating under federal and state scientific collecting permits and, for game species, in compliance with federal and state hunting regulations. Small mammals were trapped overnight using live-capture Sherman traps, following standard protocols for museum collection (Yates et al., 1996) and under state scientific collecting permits. Field research protocols were approved by the University of New Mexico Institutional Animal Care and Use Committee (Protocols 21-201225-MC and 19-200908-MC).

After collection, animals were prepared as museum specimens with an associated suite of tissues permanently frozen under ultra-cold conditions (−80°C freezers or -196°C vapor-phase liquid nitrogen storage) in the Division of Genomic Resources of the Museum of Southwestern Biology. Specimen records include spatial, temporal and natural history metadata for each sample including georeferenced locality, collection date, and reproductive and mensural data, all of which is available online through the Arctos collection management system (https://arctos.database.museum/).

All mammals were screened for ecto- and endoparasites that were also preserved and linked to mammal voucher specimens (Galbreath et al., 2019). Links to specimen records are included in Table 1. All contemporary tissues were sampled following strict protocols to avoid PFAS contamination during handling. We stored samples in PFAS-free, non-leaching polypropylene tubes (Greiner Bio-One®). Dissecting equipment was free of PFAS-containing materials and was cleaned thoroughly between specimens in HDPE (high density polyethylene) containers with Liquinox® and rinsed with deionized PFAS-free (ASTM Type II) water to avoid cross-sample contamination. Historical samples (1994) were flash frozen in liquid N_2_ and archived in NUNC® polypropylene cryovials at -80°C for ∼30 years, although specific field protocols for tissue collection were not recorded. Bird specimens were frozen for a period at -20°C before being thawed for specimen preparation, at which time tissues were dissected out and re-frozen.

We collected two plant samples, each comprised of a composite of stems and leaves from multiple individuals of an abundant shrub, four-winged saltbush, along the Holloman Lake shoreline. We screened these samples as a preliminary test of plant PFAS content and description of plant PFAS profiles to gain insights into PFAS movement from groundwater to plants to terrestrial herbivores.

### 2.2. Tissue selection for screening

We chose liver because it has been shown to be a target tissue for PFAS accumulation in multiple vertebrate classes (D’Hollander et al., 2014; Groffen et al., 2017), likely because PFAS readily bind to fatty acid binding proteins that are abundant in liver (Cara et al., 2022). We additionally chose to screen pectoral muscle for a set of game bird species because it is preferred for human consumption, although many hunters also prepare and eat liver. For a subset of the small mammals, we elected to supplement liver screening with whole blood screening. The median serum or plasma to whole blood ratio of PFOS, PFOA, PFHxS, PFNA and PFUnDA tends to be approximately 2:1; however, the ratio can vary in other PFAS, indicating that it is preferable to screen whole blood rather than serum or plasma to assay these pollutants (EFSA CONTAM Panel et al., 2020).

### 2.3. PFAS extraction and cleanup

Targeted screening for 17 PFAS substances, plus six distinct isomeric forms (Table 2) was conducted by Eurofins (Sacramento, California, USA). Before extraction, one gram of each sample was weighed out to three significant figures, thawed, and homogenized. Blood samples were gently mixed prior to aliquoting 60 µL into polypropylene centrifuge tubes, and extracted using a serial sonication with organic solvent. Known amounts of isotopically labeled Extracted Internal Standards (EISs) were immediately added to the centrifuge tube, spiked directly onto the sample. A 100 µL aliquot of ethanol and 300 µL of acetonitrile were added to each sample and briefly shaken and vortexed. The ethanol was added for analyst safety while the acetonitrile denatures blood proteins, liberating PFAS in the process. Samples were then sonicated for 20 minutes and then transferred to a refrigerated centrifuge and spun at 4700 RCF for 5 minutes. The supernatant was transferred to a new polypropylene tube. Methanol (300 µL) was added to the original tube, shaken, and vortexed before another round of centrifugation. Supernatants were combined and diluted to <10% organic prior to loading onto pre-conditioned weak anion exchange (WAX) solid phase extraction (SPE) cartridges (30 mg/3 mL). Samples were eluted with a small volume of basic methanol, the non-extracted internal standard (NIS) added to the extract and the final volume was adjusted to 240 µL with water to achieve 80:20 methanol:water composition for analysis.

**Table 2.**
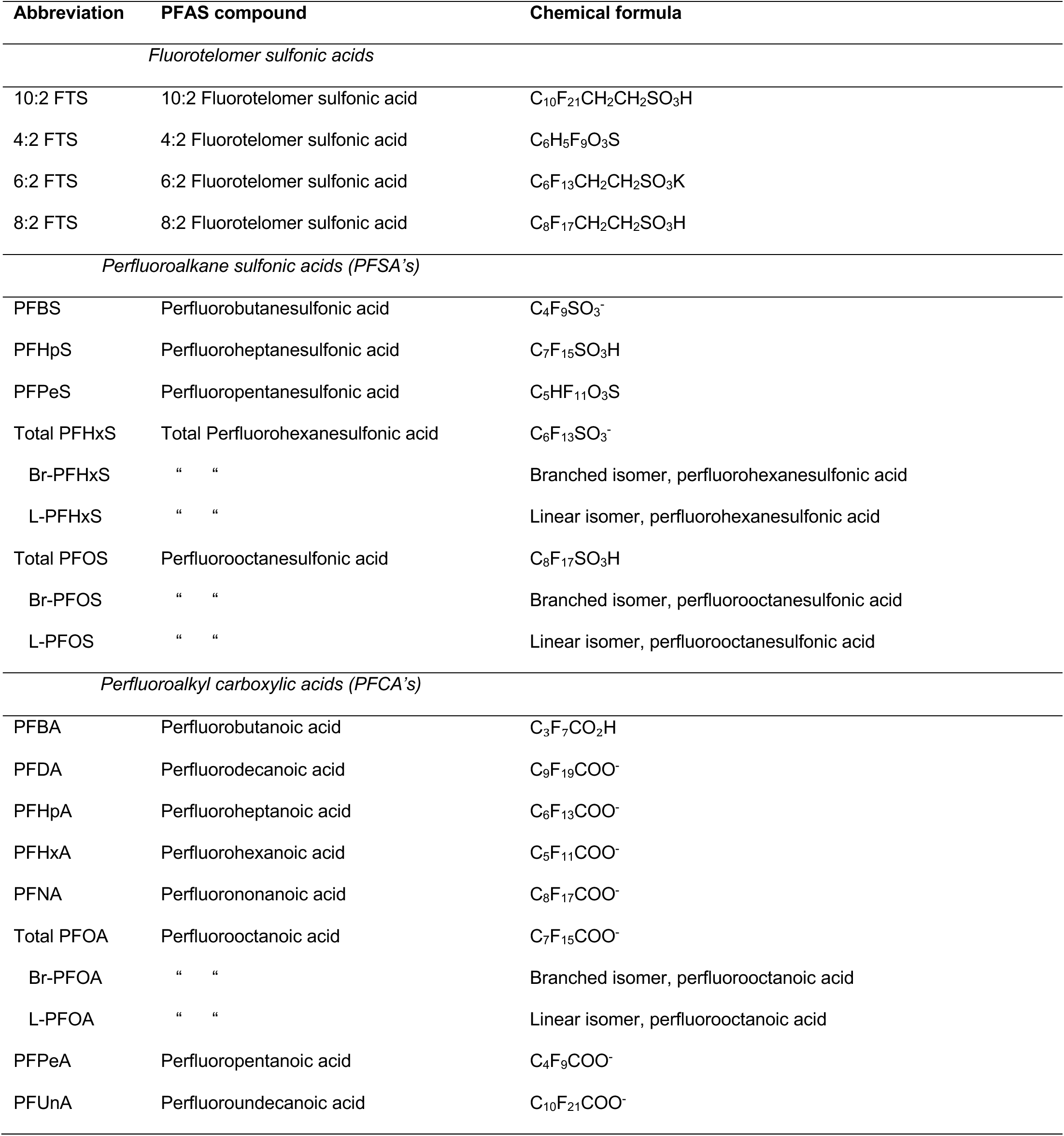
List of 17 PFAS compounds and 6 alternative isomers assayed in this study. PFSA’s with 6 or more C atoms and PFCA’s with 7 or more C atoms are considered ‘long-chain’ (Buck et al., 2011).

| Abbreviation | PFAS compound | Chemical formula |
| --- | --- | --- |
| *Fluorotelomer sulfonic acids* |  |  |
| 10:2 FTS | 10:2 Fluorotelomer sulfonic acid | $C_{10}F_{21}CH_2CH_2SO_3H$ |
| 4:2 FTS | 4:2 Fluorotelomer sulfonic acid | $C_6H_5F_9O_3S$ |
| 6:2 FTS | 6:2 Fluorotelomer sulfonic acid | $C_6F_{13}CH_2CH_2SO_3K$ |
| 8:2 FTS | 8:2 Fluorotelomer sulfonic acid | $C_8F_{17}CH_2CH_2SO_3H$ |
| *Perfluoroalkane sulfonic acids (PFSA's)* |  |  |
| PFBS | Perfluorobutanesulfonic acid | $C_4F_9SO_3^-$ |
| PFHpS | Perfluoroheptanesulfonic acid | $C_7F_{15}SO_3H$ |
| PFPeS | Perfluoropentanesulfonic acid | $C_5HF_{11}O_3S$ |
| Total PFHxS | Total Perfluorohexanesulfonic acid | $C_6F_{13}SO_3^-$ |
| Br-PFHxS | " " | Branched isomer, perfluorohexanesulfonic acid |
| L-PFHxS | " " | Linear isomer, perfluorohexanesulfonic acid |
| Total PFOS | Perfluorooctanesulfonic acid | $C_8F_{17}SO_3H$ |
| Br-PFOS | " " | Branched isomer, perfluorooctanesulfonic acid |
| L-PFOS | " " | Linear isomer, perfluorooctanesulfonic acid |
| *Perfluoroalkyl carboxylic acids (PFCA's)* |  |  |
| PFBA | Perfluorobutanoic acid | $C_3F_7CO_2H$ |
| PFDA | Perfluorodecanoic acid | $C_9F_{19}COO^-$ |
| PFHpA | Perfluoroheptanoic acid | $C_6F_{13}COO^-$ |
| PFHxA | Perfluorohexanoic acid | $C_5F_{11}COO^-$ |
| PFNA | Perfluorononanoic acid | $C_8F_{17}COO^-$ |
| Total PFOA | Perfluorooctanoic acid | $C_7F_{15}COO^-$ |
| Br-PFOA | " " | Branched isomer, perfluorooctanoic acid |
| L-PFOA | " " | Linear isomer, perfluorooctanoic acid |
| PFPeA | Perfluoropentanoic acid | $C_4F_9COO^-$ |
| PFUnA | Perfluoroundecanoic acid | $C_{10}F_{21}COO^-$ |

Thawed, homogenized tissues for extraction were spiked with EIS then solvated with 0.4% KOH in methanol and sonicated for one hour. Samples were separated by centrifuge at 4700 RCF followed by decanting the supernate to a new vessel prior to a second 0.4% KOH/methanol solvation. This solution was shaken briefly and sonicated and separated as above. Supernates were combined and the sample diluted to 125 mL with deionized water and neutralized with glacial acetic acid to pH 6–8. This extract was then passed over a conditioned WAX/graphite carbon black (GCB) SPE (500 mg, 50 mg, 6 mL). Cartridges were washed with small volumes of methanol in water prior to drying and then eluted with 0.3% ammonium hydroxide in methanol. Samples were not concentrated after SPE, but rather spiked with NIS and brought to 10 mL of 80:20 methanol:water prior to analysis.

One Method Blank (MB) and two Laboratory Control Samples (LCS/LCS Duplicate) containing 2% bovine serum albumin (BSA) for blood batches and 0.02 g corn oil for tissue batches were included with each field sample at a rate of 5% (MB, LCS/LCSD per 20 samples). The LCS/LCSD were also spiked with known concentrations of native PFAS. These QC samples are standard practice among US EPA methods and serve as both positive (LCS/LCSD) and negative (MB) controls as they follow all field samples throughout extraction and analysis.

### 2.4. Analysis by mass spectrometry

The mass spectrometer analytical configuration consisted of an Exion LC system with two pumps and a SCIEX 7500 for blood analyses and a SCIEX 5500 for tissue analyses. Pump-A contained 20 mM ammonium acetate and pump-B delivered 0.5% NH_4_OH in methanol. Pump-A was used to load 20 µL of sample extract onto an analytical column (Gemini 3 µm C18 50X2 mm) heated to 45° C where compounds of interest were separated before introduction into the instrument source. The electrospray source was operated in negative polarity and data acquired in multiple reaction monitoring (MRM) mode with a minimum of 10 scans per peak. Prior to analysis, optimal voltages for both ionization and collision induced disassociation were determined by direct infusion. Analysis time for this process resulted in an approximate 9-minute cycle time. See Supplemental Data 1–6 for additional details on PFAS assays.

### 2.5. Data analysis

All assayed concentrations of individual PFAS and isomers were summarized, compared, and visualized in R. To assess differences among contamination profiles of species, tissues, localities, and substances, we created a heat map with log_10_-transformed PFAS concentrations and applied marginal dendrograms to visualize similarity among species-tissue-locality combinations and substances, respectively. We conducted classical non-metric multidimensional scaling (NMDS) to assess variation among individual animal tissues with respect to their PFAS contamination profiles and in relation to sampling site (Holloman AFB vs. control). Because sample sizes for species-tissue combinations from control localities were low (n ≤ 9), leading to unreliable assessments of normality, we tested for differences between localities with non-parametric tests (Altman et al., 1983). For each assayed PFAS, we conducted non-parametric Kruskal-Wallis tests to compare birds and small mammals from the contaminated Holloman AFB sites with those from control sites.

We compared concentrations among tissues within individuals to quantify tissue-specific exposure, toxic potency, and bioaccumulation (Gomis et al., 2018). We used paired t-tests and F-tests to assess equal variance between bird tissues from contaminated sites, where sampling was adequate for parametric approaches. Due to unequal variances between tissue types for all PFAS compounds, we used paired-sample Mann-Whitney Wilcoxon Signed Rank tests with a normal approximation to compare PFAS concentrations in bird muscle tissue to liver tissue and mammal blood to liver tissue from the contaminated sites. We note that tissue comparisons should be interpreted with caution because of the possibility of matrix effects, such as ion enhancement or suppression that can add systematic error (Berger and Haukås, 2005).

## 3. Results

PFAS were strikingly abundant and widespread among animal tissues that we sampled at Holloman AFB. We obtained measurements above reporting limits for 16 of 17 PFAS (Table S1), and the most common PFAS were detected above reporting limits in the overwhelming majority of Holloman AFB vertebrate samples (Table 3, S1). For Holloman liver samples, mammals had higher mean concentrations of total PFAS (ΣPFAS; x̄ = 15,589 ng/g, SD = 24,623) than birds (x̄ = 11,508 ng/g, SD = 11,284), although this difference was driven largely by ΣPFAS high concentrations in house mice (*Mus musculus*) (x̄ = 28,276 ng/g, SD = 25,644); mammals excluding house mice averaged lower (x̄ = 8,173 ng/g, SD = 21,062).

**Table 3.**
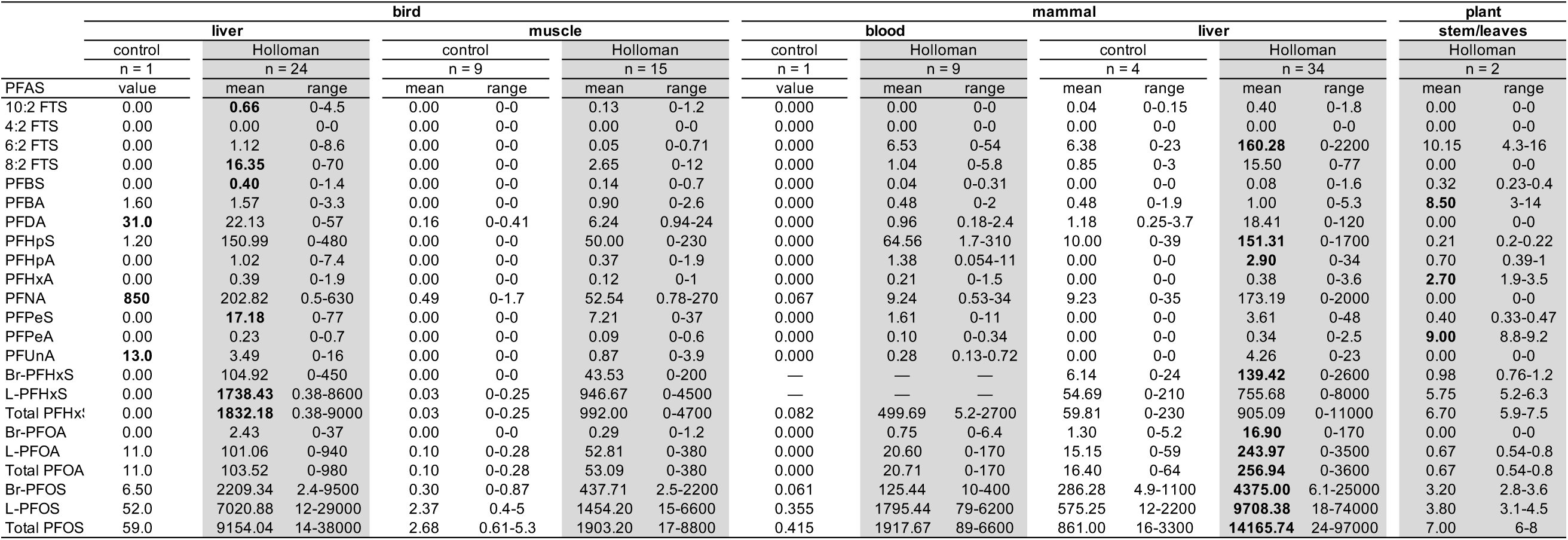
Detection levels for 17 PFAS compounds and six isomeric constituents in bird liver (ng/g), bird pectoral muscle (ng/g), mammal blood (ng/mL), mammal liver (ng/g), and plant stem & leaf tissue (ng/g), respectively, for Holloman AFB and control sites. Bold values represent the highest mean values for each compound across the nine tissue sample types.

Vertebrate sample types, when grouped by species, tissue type, and locality, fell into two large clusters with respect to PFAS composition and concentrations (Fig. 1). The first group comprised only Holloman AFB samples that were highly contaminated with a broad range of PFAS, including all of the aquatic-habitat birds except mergansers, as well as house mice and four other rodent species (Fig. 1). The second group comprised the remaining rodent species and an upland desert songbird species from Holloman AFB as well as all of the control-site samples, (Fig. 1). We found multiple PFAS in the tissues of all eleven game-bird species that were tested at Holloman AFB, ten out of eleven species contained high levels. Only the common mergansers (n = 2) collected at Holloman Lake were not highly contaminated and more closely resembled control samples in their PFAS profiles (Fig. 1).

PFAS in animal tissues was strikingly higher at Holloman AFB than at control sites (Table 3, Figs. 1, 2, S1). Among all taxa and tissues, samples from Holloman AFB were more than 30-fold higher in ΣPFAS (x̄ = 10,527 ng/g or ng/mL, SD = 17,641) than samples from control sites (x̄ = 324 ng/g or ng/mL, SD = 965). Among bird muscle samples, 13 of 17 targeted PFAS substances (plus six constituent isomers) were detected at significantly higher levels in Holloman AFB samples relative to control sites: PFBA, PFHpA, L-PFOA, Br-PFOA, Total PFOA, PFNA, PFDA, PFUnA, PFBS, PFPeS, Br-PFHxS, L-PFHxS, Total PFHxS, PFHpS, L-PFOS, Br-PFOS, Total PFOS, 8:2 FTS, and 10:2 FTS (Fig. 2; full names of each compound listed in Table 2). Among mammal liver samples, four PFAS and two constituent isomers were measured at significantly higher levels among Holloman AFB samples relative to control sites (despite that statistical power was limited by the small number of control samples assayed): PFDA, PFUnA, PFHpS, L-PFOS, Br-PFOS, and Total PFOS (Fig. S1).

**Fig. 2.**
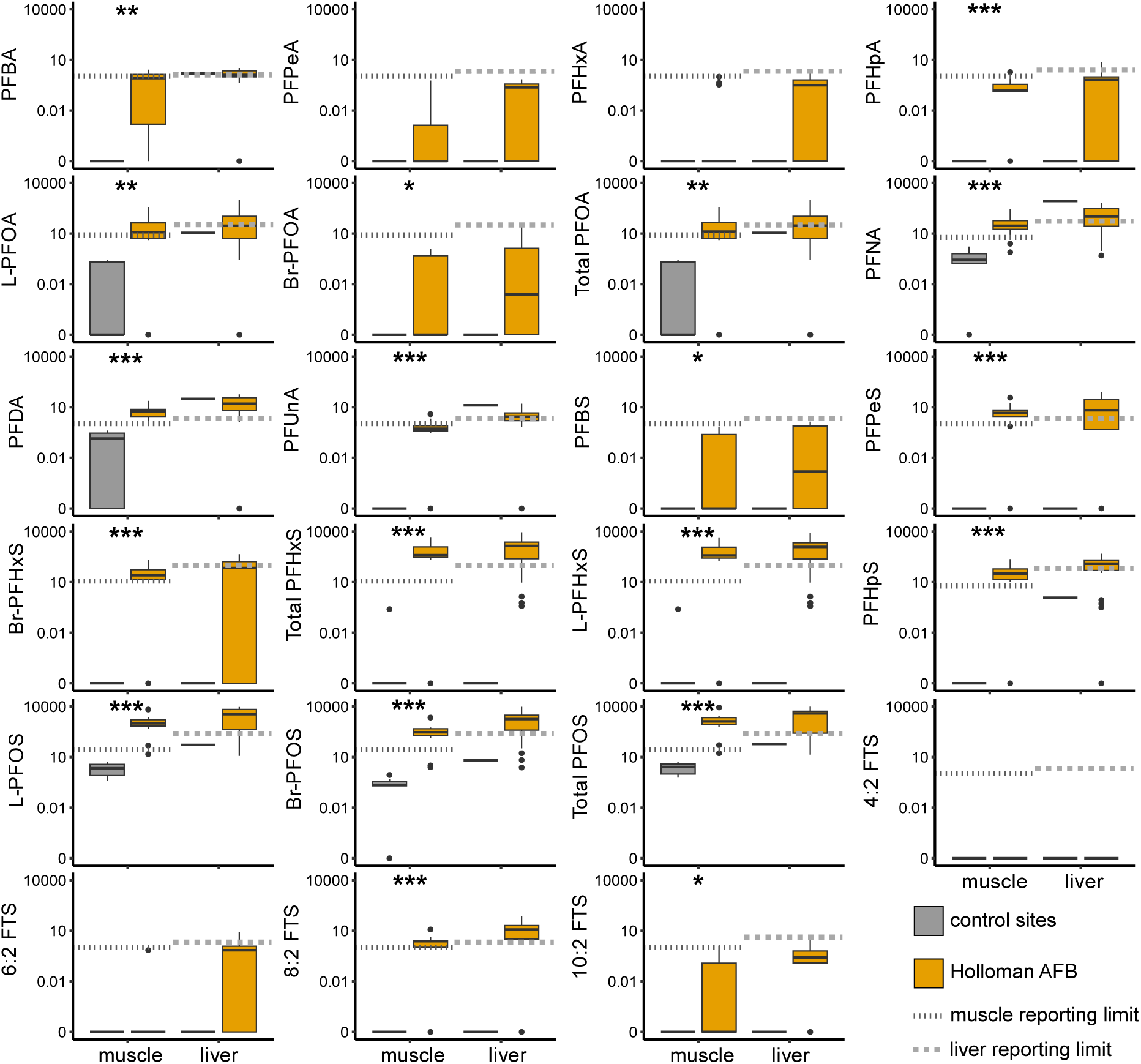
Comparisons of PFAS concentrations (ng/g) in bird muscle and liver samples from Holloman AFB (orange) versus offsite control localities (gray). Gray dotted lines and dashed lines indicate mean reporting limit thresholds per compound for muscle and liver, respectively. Asterisks indicate significant differences, calculated from Kruskal-Wallis tests (for muscle samples only; *p< 0.05; **p < 0.005; ***p < 0.0005). Because we had only a single liver sample from a control site, we did not run significance tests for liver. The liver data from the one control site sample, a northern pintail, had low concentrations of PFOS, PFOA, and PFHxS. However, this sample was unique in showing concentrations well above the reporting limit for three other compounds (PFNA, PFDA, PFUnA), suggesting that this duck had become contaminated at a site with a different profile of PFAS contaminants during the course of its migratory cycle. No muscle samples from control sites showed elevated PFAS concentrations or anomalous PFAS profiles.

PFOS was the most abundant PFAS in animal tissues, both at Holloman AFB and control sites, followed by PFHxS (Table 3, Figs. 3 & S1). A white-footed mouse (*Peromyscus leucopus*) collected in 1994 had the highest PFAS level among all samples (liver [PFOS] = 97,000 ng/g ww); however, the four samples from 1994 exhibited a wide range of PFOS levels (as low as 24 ng/g ww) and could not be distinguished from modern samples based on their PFAS profiles. An American wigeon (*Mareca americana*) collected in 2022 had the highest PFAS levels among sampled birds (liver [PFOS] = 38,000 ng/g ww). 4:2 fluorotelomer sulfonic acid (FTS) was the only tested substance that we never detected above reporting limits (Table 3).

**Fig. 3.**
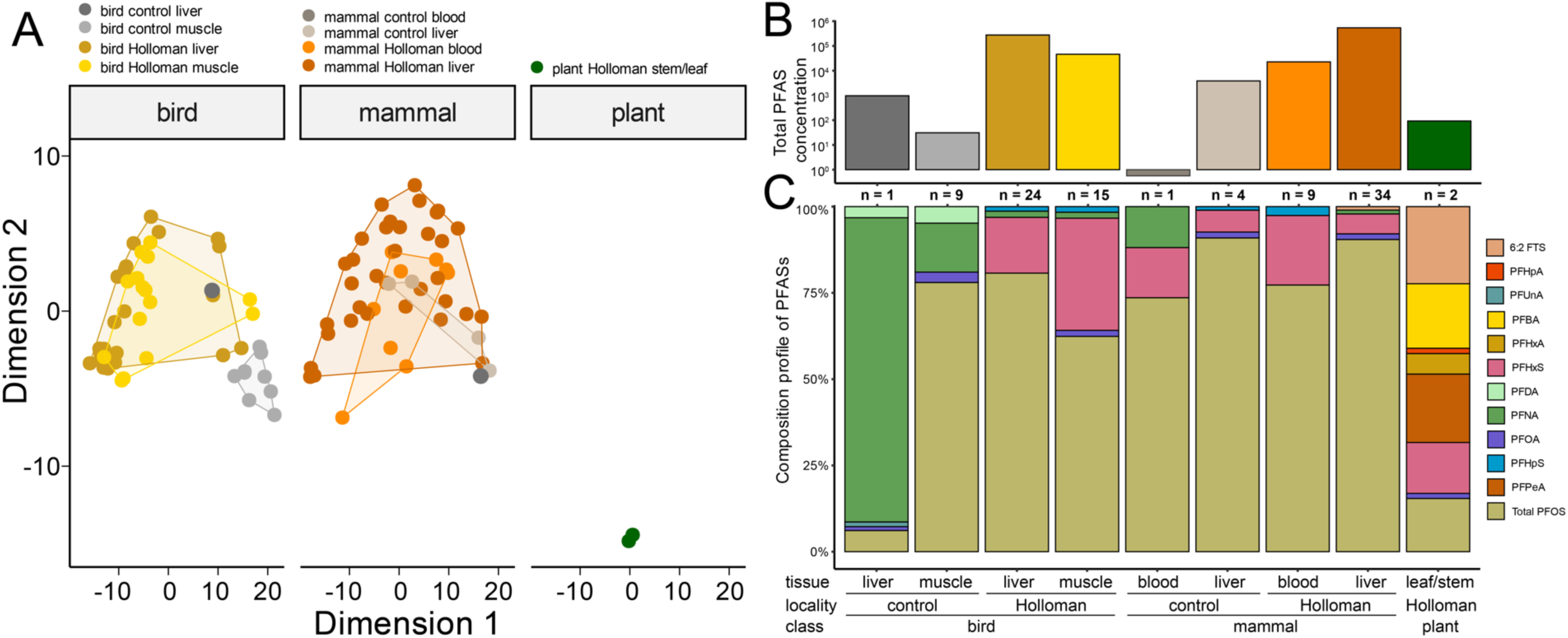
(A) Multidimensional scaling plot showing locality-tissue clusters, separated by clade for clarity, of PFAS concentrations from Holloman Air Force Base and offsite control localities. Clusters are bounded by convex hulls. (B) ΣPFAS concentrations (ng/g, or ng/ml for blood) colored by clade-locality-tissue combination (see key in panel A). (C) Composition of PFAS that comprised at minimum 1% of total contaminants, averaged by clade and tissue from control sites and Holloman Lake, respectively. X-axis labels are shared between panels B and C.

Overall, there were strong similarities of the PFAS contamination profiles among bird and mammal samples (Figs. 1, 3). Most bird and mammal tissues from Holloman AFB sites clustered along the first two NMDS dimensions (Fig. 3A). Control samples including bird muscle samples and a single mammal blood sample clustered separately (Fig. 3A). PFAS composition was dominated by PFOS (Fig. 3B), averaging ∼10,000 ng/g ww in liver for both birds and mammals from Holloman AFB sites. One prominent exception was a northern pintail liver sample from a control site that contained the highest levels of PFNA of any sample in our study (850 ng/g ww) but very little PFOS. PFHxS was the second most abundant PFAS detected, averaging in the 1000s of ng/g ww among bird and mammal tissues from Holloman AFB. Together PFOS and PFHxS comprised >75% of ΣPFAS. Additional abundant PFAS in vertebrate tissues at Holloman AFB included PFOA, PFNA, PFHpS, 6:2 FTS, PFDA, 8:2 FTS, and PFPeS (Fig. 3C).

Two key differences between birds and mammals emerged from our data. First, PFHxS and PFOA were far more abundant in avian liver than in mammal liver, with the exception that the house mice tended to resemble the birds (Fig. 1). Second, 6:2 FTS was found at high levels in some mammal livers (x̄ =160.3 ng/g ww), but not avian livers (x̄ =1.1 ng/g ww) (Table 3). Samples of the plant species (four-wing saltbush) from Holloman Lake clustered apart from the animals, with lower overall levels of PFAS and starkly contrasting composition (6:2 FTS was the most abundant compound; Fig. 3).

For both birds and mammals, liver showed evidence of higher bioaccumulation than other tissues, as expected (Figs. 4 & S2). Eleven out of 17 PFAS were significantly higher in bird liver than muscle (Fig. 4). Mammals showed higher bioaccumulation in liver than blood; seven out of 17 PFAS were significantly more concentrated in mammal liver tissue than blood from the same individuals (Fig. S2).

**Fig. 4.**
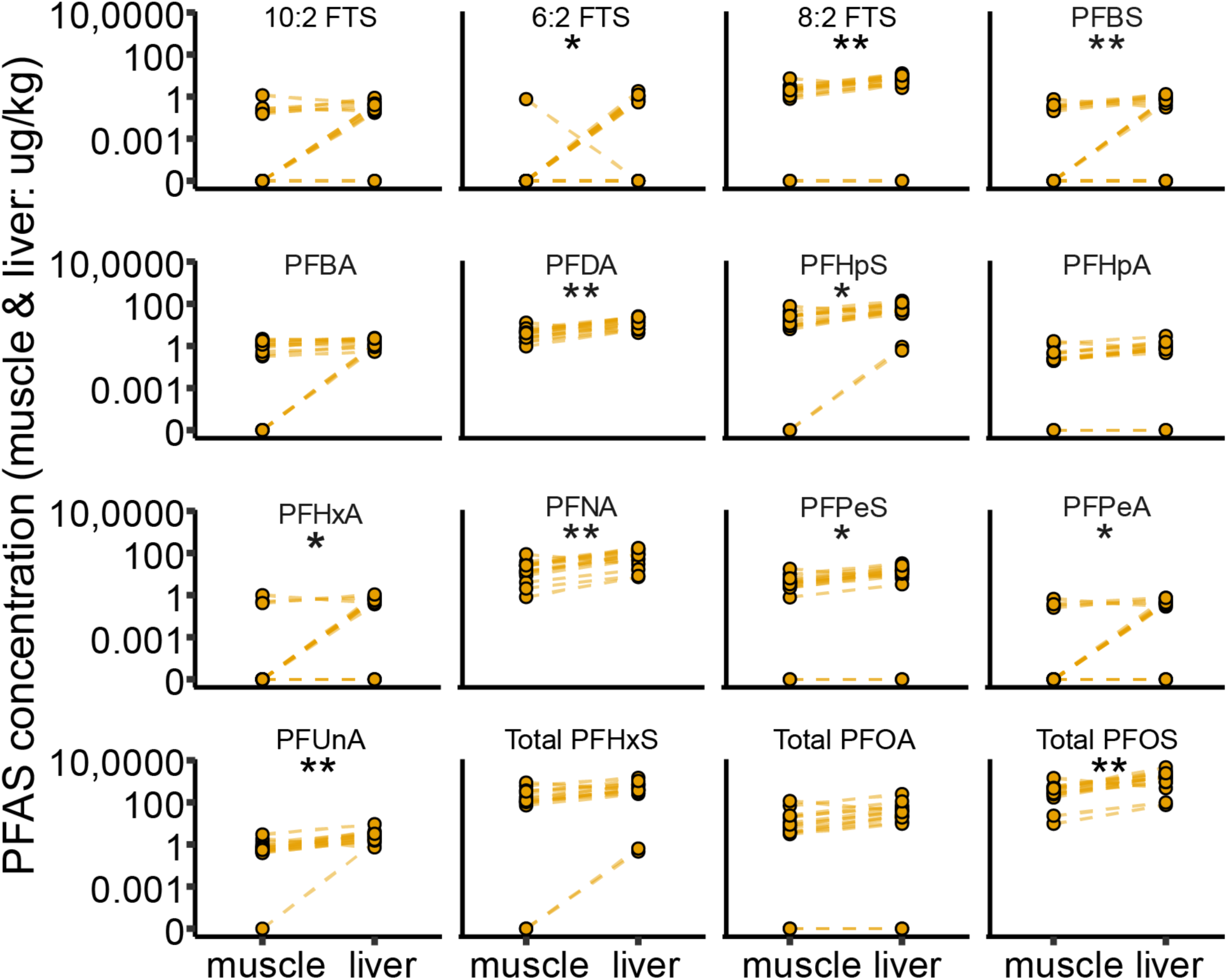
Between-tissue comparison for muscle and liver of birds. Points indicate PFAS concentrations for muscle or liver, measured in ng/g. Lines connect points from the same animal (n = 30). Steeper lines connecting tissues within individuals represent larger between tissue differences in PFAS concentrations. Asterisks indicate significant differences in PFAS concentrations between tissues based on Wilcoxon Signed Rank tests using normal approximation (*p < 0.05; **p < 0.005)

We separately assayed linear (L-) and branched (Br-) isomers for three PFAS: PFOA, PFHxs, and PFOS (Fig. 4). All tissues in all taxa were substantially enriched for L-PFOA, with a mean proportion of 0.98 ± 0.03%, ∼20% higher than typical manufactured PFOA (Fig. S3, A–B) (Buck et al., 2011). All tissues and taxa were similarly enriched in L-PFHxS relative to Br-PFHxS (μ = 0.93 ± 0.08%), though mammal liver showed significant variation in the proportion of L-PFHxS (0.49 – 0.90%) (Fig. S3, C). Though variable among tissues and taxa, the proportion of L-PFOS in all taxa and tissues (μ = 0.75 ± 0.09%; range = 0.52 – 0.95%) fell near the typical manufacturing proportion of 70% L-PFOS (Fig. S3). Relative to manufacturing standards, mammalian blood showed the highest enrichment in L-PFOS (μ = 0.93 ± 0.02%), while the plant (four-wing saltbush) tissue was depleted in L-PFOS (μ = 0.54 ± 0.03%) (Fig. S3, E & F).

## 4. Discussion

### 4.1. A contaminated desert oasis

Diverse wild animal species at contaminated wetlands and upland sites on and adjacent to the Holloman AFB harbor high levels of PFAS in their tissues; with very high levels of PFAS contamination from at least 1994 to the present. PFAS have permeated the food web at concentrations that are concerning for livestock, wildlife, and human health. Wild bird and rodent species are potentially exposed to high PFAS levels through their diet as well as through contact, inhalation, or incidental ingestion of environmental PFAS. We provide the first direct evidence that PFAS originating on a New Mexico Air Force Base actively contaminate the thousands of migratory birds that use these wetlands as breeding, wintering, or migratory stopover habitats. Specific PFAS that were most abundant in animal tissues at Holloman Lake were predictable based on previous studies of water and sediment at U.S. military installations contaminated with AFFFs (Anderson et al. 2016). Accordingly, PFOS and PFHxS were the most abundant PFAS detected. PFOA and a series of other compounds were also present at high levels, but not in all samples.

### 4.2. Why PFAS concentrations at Holloman AFB should be considered extraordinary

The birds and rodents at Holloman AFB had PFAS tissue concentrations far higher than those measured in nearly all previous surveys of wildlife, with liver concentrations of PFOS and ΣPFAS, respectively, averaging in the tens of thousands of ng/g ww. For perspective, at Cannon AFB in eastern New Mexico, ground water contamination with PFAS led to the contamination and subsequent destruction of ∼5000 dairy cattle; their milk contained PFOS in the hundreds to thousands of parts per trillion, 3–4 orders of magnitude *lower* than the tissue concentrations that we measured in Holloman wildlife (Jha et al., 2021).

The highest previous measurements in wild populations were from a point source near Antwerp, Belgium, where a 3M fluorochemical plant contaminated the environment, including local vertebrate species such as wood mice (*Apodemus sylvaticus*) and great tits (*Parus major*; a songbird). Wood mouse livers collected near the plant in 2002 contained a median of ∼5,000 ng/g ww, but one sample contained an astounding 179,000 ng/g ww, while two others measured 98,000 ng/g ww (Hoff et al., 2004)––similar to the highest level of contamination that we observed at Holloman AFB. Two years later, in 2004, the highest measurement from a wood mouse liver at the site was ∼22,000 ng/g ww (D’Hollander et al., 2014). For great tit eggs within 1 km of the plant in 2011, half of the samples harbored PFOS in the 10,000s of ng/g ww, and the highest measured level at 69,218 ng/g ww (Groffen et al., 2017); liver PFOS in great tits at the same site was measured as high as 11,359 ng/g ww (Dauwe et al., 2007).

Away from point sources, PFAS tissue concentrations tend to be lower, but with occasional, heavily contaminated outliers. Giesy and Kannon reported PFAS from wildlife around the world and only bald eagle (*Haliaeetus leucocephalus*) and mink (*Neogale vison*) were as high as the 1000s of ng/g ww, with a river otter (*Lontra canadensis*) reaching 990 ng/g ww (Giesy and Kannan, 2001). Subsequently, Eurasian otters (*Lutra lutra*) in northern Europe (Androulakakis et al., 2022), river otters in Illinois, USA (Kannan et al., 2002), and minks in Sweden (Persson and Magnusson, 2015) were reported as high as the thousands of ng/g ww. In a 2007 review of wildlife screening data for 46 species, 29 had maximum measured tissue concentrations in the tens of ng/g ww or below, and only two reached the 1000s of ng/g ww: great tit from a contaminated point source reached 1625 ng/g ww, and bald eagles in North America ranged as high as 2570 ng/g ww (Lau et al., 2007). At high latitudes in the North Atlantic and southern Indian oceans, PFAS have been detected in marine mammal and seabird tissues and eggs since at least the 1970s (Kelly et al., 2009; Bianchini et al., 2022) and typically occurs in the single-digits to 100’s of ng/g ww, occasionally ranging to the 1000s of ng/g ww (Verreault et al., 2005; Taylor et al., 2021). By contrast, seven marine and terrestrial mammals important for subsistence hunting on Sivuqaq Island (St. Lawrence Island), Alaska, harbored very low PFAS (Byrne et al., 2022); although one of the Sivuqaq study species, reindeer (*Rangifer tarandus*), was found to have liver ΣPFAS concentrations >100 ng/g in Greenland (Roos et al., 2022). A suite of wild avian scavenger and predator species, which are known to range widely, have been found to harbor moderate blood and tissue concentrations of PFOS, typically in the singles to hundreds of ng/g ww, with one carrion crow in Japan measuring to 1200 ng/g ww (Taniyasu et al., 2003), and only bald eagle having been repeatedly found to range to the low 1000s of ng/g ww (Verreault et al., 2005; Lopez-Antia et al., 2021; Herzke et al., 2023; Badry et al., 2022). In China, shorebird (Charadriiformes) eggs averaged in the tens of ng/g for PFOS and PFOA, with PFOA reaching a high of 121 ng/g ww (Sun et al., 2023). Among bird liver and eggs for a suite of species in South Korea, PFOS comprised ∼80% of ΣPFAS, most species ranged to the hundreds of ng/g, seven had liver concentrations of ΣPFAS ranging into the thousands of ng/g, and one outlier individual, a scops owl (*Otus* sp.), had 11,283 ng/g ww (Yoo et al., 2008; Barghi et al., 2018; Park et al., 2021). Japanese field mice (*Apodemus speciosus*) at contaminated and non-contaminated sites from across Japan averaged 5.7 ng/ml PFOS with a maximum of 148 ng/ml (Taniyasu et al., 2013). Taken together, the previously published surveys of PFAS concentrations in wildlife show that the levels of PFAS contamination that we found in animal tissues at Holloman AFB were highly unusual, as was the degree to which PFAS has permeated the community of secondary consumers.

### 4.3. Effects on animal health

Toxicity reference values for birds are few and variable, but the available toxicity data suggest that the avian liver concentrations of PFOS that we recorded at Holloman AFB are sufficient to damage health and diminish ecological performance. Newsted et al. estimated ‘predicted no effect concentrations’ (PNEC; 350 ng/g ww) and a toxicity reference value for upper-trophic avian predators (TRV; 600 ng/g ww) (Newsted et al., 2005). Dennis et al. reported chronic toxicity reference values (CRVs) of PFOS as 226 and 50.4 ng/g ww liver tissue for adults and juveniles, respectively, of northern bobwhites (*Colinus virginianus*) that were chronically exposed to PFOS or a simple PFAS mixture via drinking water; these latter thresholds may be most relevant with respect to chronically exposed birds at Holloman AFB (Dennis et al., 2021). Dennis et al. further observed that PFOS was absorbed and distributed differently when combined with PFHxS (as it is at Holloman AFB) and these interactive effects may help to explain the variability in outcomes of toxicity studies. Every tissue sample of a game bird species from Holloman AFB exceeded the CRV threshold of 226 ng/g ww, except for three of the four common merganser samples (Table 4).

**Table 4.** Total PFOS, PFOA, and PFHxS, respectively, by species and tissue for Holloman AFB and control samples. Units are ng/g except for blood for which they are ng/mL.

| Locality | Class | English name | Tissue type | n | Mean total PFOS | PFOS range | Mean total PFOA | PFOA range | mean total PFHxS | PFHxS range |
| --- | --- | --- | --- | --- | --- | --- | --- | --- | --- | --- |
| Holloman | bird | American Coot | liver | 1 | 5000.0 | - | 17.0 | - | 1200.0 | - |
| control | bird | American Coot | muscle | 5 | 3.0 | 0.77–5.3 | 0.1 | 0–0.28 | 0.1 | 0–0.25 |
| Holloman | bird | American Coot | muscle | 1 | 960.0 | - | 4.1 | - | 330.0 | - |
| Holloman | bird | American Wigeon | liver | 1 | 38000.0 | - | 40.0 | - | 3500.0 | - |
| Holloman | bird | American Wigeon | muscle | 1 | 2100.0 | - | 5.2 | - | 500.0 | - |
| Holloman | bird | Bufflehead | liver | 1 | 20000.0 | - | 980.0 | - | 9000.0 | - |
| Holloman | bird | Bufflehead | muscle | 1 | 2700.0 | - | 200.0 | - | 3000.0 | - |
| Holloman | bird | Common Goldeneye | liver | 1 | 2200.0 | - | 100.0 | - | 1700.0 | - |
| Holloman | bird | Common Goldeneye | muscle | 1 | 8800.0 | - | 380.0 | - | 4700.0 | - |
| Holloman | bird | Common Merganser | liver | 2 | 265.0 | 220–310 | 0.0 | 0–0 | 0.5 | 0.38–0.57 |
| Holloman | bird | Common Merganser | muscle | 2 | 34.0 | 17–51 | 0.0 | 0–0 | 0.0 | 0–0 |
| Holloman | bird | Common Yellowthroat | liver | 2 | 5750.0 | 4300–7200 | 9.2 | 7.3–11 | 285.0 | 280–290 |
| Holloman | bird | Green-winged Teal | liver | 3 | 14333.3 | 11000–20000 | 66.3 | 47–97 | 2266.7 | 1600–3000 |
| control | bird | Green-winged Teal | muscle | 1 | 1.0 | - | 0.0 | - | 0.0 | - |
| Holloman | bird | Green-winged Teal | muscle | 3 | 1536.7 | 810–2600 | 9.4 | 5.6–13 | 413.3 | 280–640 |
| Holloman | bird | Horned Lark | liver | 2 | 76.5 | 23–130 | 0.9 | 0–1.7 | 5.0 | 0.6–9.4 |
| Holloman | bird | Killdeer | liver | 2 | 14200.0 | 5400–23000 | 77.5 | 15–140 | 1095.0 | 390–1800 |
| control | bird | Mallard | muscle | 2 | 3.9 | 2.6–5.1 | 0.0 | 0–0 | 0.0 | 0–0 |
| control | bird | Northern Pintail | liver | 1 | 59.0 | - | 11.0 | - | 0.0 | - |
| Holloman | bird | Northern Pintail | liver | 1 | 10000.0 | - | 22.0 | - | 1800.0 | - |
| Holloman | bird | Northern Pintail | muscle | 1 | 1300.0 | - | 4.9 | - | 400.0 | - |
| Holloman | bird | Northern Shoveler | liver | 3 | 7371.3 | 14–17000 | 147.8 | 0.27–360 | 2500.5 | 1.4–5900 |
| Holloman | bird | Northern Shoveler | muscle | 2 | 1600.0 | 1100–2100 | 33.5 | 15–52 | 900.0 | 400–1400 |
| Holloman | bird | Redhead | liver | 2 | 11850.0 | 9700–14000 | 160.0 | 120–200 | 2650.0 | 1000–4300 |
| Holloman | bird | Redhead | muscle | 2 | 1255.0 | 610–1900 | 30.5 | 20–41 | 655.0 | 210–1100 |
| Holloman | bird | Ruddy Duck | liver | 1 | 9400.0 | - | 170.0 | - | 3800.0 | - |
| control | bird | Ruddy Duck | muscle | 1 | 0.6 | - | 0.2 | - | 0.0 | - |
| Holloman | bird | Ruddy Duck | muscle | 1 | 2300.0 | - | 46.0 | - | 1600.0 | - |
| Holloman | bird | Song Sparrow | liver | 2 | 2850.0 | 1800–3900 | 9.1 | 1.2–17 | 300.0 | 170–430 |
| Holloman | dicot | Four-wing saltbush | leaf & stem | 2 | 7.0 | 6–8 | 0.7 | 0.54–0.8 | 6.7 | 5.9–7.5 |
| Holloman | mammal | Cactus mouse | liver | 1 | 22000.0 | - | 3.3 | - | 38.0 | - |
| control | mammal | Chihuahuan pocket mouse | liver | 2 | 1664.0 | 28–3300 | 32.0 | 0–64 | 115.3 | 0.64–230 |
| Holloman | mammal | Chihuahuan pocket mouse | liver | 2 | 4420.0 | 440–8400 | 25.6 | 4.2–47 | 357.0 | 34–680 |
| Holloman | mammal | Desert pocket gopher | blood | 1 | 89.0 | - | 0.0 | - | 12.0 | - |
| Holloman | mammal | Hispid cotton rat | blood | 2 | 370.0 | 310–430 | 0.1 | 0–0.27 | 170.0 | 110–230 |
| Holloman | mammal | Hispid cotton rat | liver | 5 | 2000.0 | 1000–3900 | 0.4 | 0–0.65 | 88.4 | 36–160 |
| Holloman | mammal | House mouse | blood | 2 | 4250.0 | 1900–6600 | 92.5 | 15–170 | 1620.0 | 540–2700 |
| Holloman | mammal | House mouse | liver | 13 | 24081.6 | 61–65000 | 666.5 | 3.1–3600 | 2244.8 | 2.2–11000 |
| Holloman | mammal | Chihuah. grasshopper mouse | blood | 1 | 5900.0 | - | 0.8 | - | 720.0 | - |
| Holloman | mammal | Chihuah. grasshopper mouse | liver | 1 | 850.0 | - | 0.0 | - | 37.0 | - |
| control | mammal | Merriam's kangaroo rat | blood | 1 | 0.4 | - | 0.0 | - | 0.1 | - |
| Holloman | mammal | Merriam's kangaroo rat | blood | 2 | 215.0 | 160–270 | 0.0 | 0–0 | 17.6 | 5.2–30 |
| control | mammal | Merriam's kangaroo rat | liver | 2 | 58.0 | 16–100 | 0.8 | 0–1.6 | 4.3 | 0–8.6 |
| Holloman | mammal | Merriam's kangaroo rat | liver | 4 | 1165.0 | 700–1700 | 0.0 | 0–0 | 8.5 | 0–27 |
| Holloman | mammal | Western Harvest Mouse | liver | 4 | 4475.0 | 1600–7500 | 0.0 | 0–0 | 14.8 | 0–33 |
| Holloman | mammal | White-footed mouse | blood | 1 | 1600.0 | - | 0.3 | - | 150.0 | - |
| Holloman | mammal | White-footed mouse | liver | 4 | 26081.0 | 24-97000 | 3.9 | 0-15 | 66.7 | 0.86-160 |

For rodents and other non-avian vertebrates, there is also a shortage of PFAS toxicity studies for wild populations. A ‘no-observed-adverse-effect-concentration’ (NOAEC) of 50,000 ng/mL has been estimated as repeated dose toxicity in rodent serum (Colnot and Dekant, 2022); but just as is the case in birds, adverse effects have been reported at far lower concentrations.

Previous work has demonstrated striking health consequences for vertebrate populations even when PFAS occurred at much lower tissue concentrations than we recorded in this study. In a study of three species of gulls (Charadriiformes) from France, declines in condition and thyroid hormone disruption were associated with several PFAS, but concentrations were in the low single ng/g ww for most compounds, and as high as the low tens of ng/g ww for PFOS (Sebastiano et al., 2023). Similar levels of PFOS were associated with lower antioxidant capacity in another gull species, the black-legged kittiwake (Costantini et al., 2019). Small doses of PFOS affected body weight and reproduction of northern bobwhites (Ankley et al., 2021). In domestic and feral cats, PFAS increased body weight and risk of liver, thyroid, and kidney disease at blood concentrations averaging 6.9 ng/ml PFHxS (maximum 235 ng/ml), 8.9 ng/ml PFOS (maximum 121 ng/ml) (Bost et al., 2016) and 9.5 ng/ml ΣPFAS (Wang et al., 2018). Police dogs exposed to firefighting foams and laboratory beagles exposed to dietary PFAS showed dose-dependent alterations in amylase, cholesterol, and several indicators of blood chemistry at ΣPFAS levels averaging 3.6 ng/ml (maximum 16.6 ng/ml) (You et al., 2022). Aquatic secondary consumers, mainly fish, were adversely affected by serum PFAS as low as 13.5 ng/mL (Banyoi et al., 2022). Immunotoxicity resulted from very low daily doses of 6:2 FTS in white-footed mice (Bohannon et al., 2023). Alligators show autoimmune-like effects from modest levels of PFAS in serum, averaging 27 ng/mL, but as high as 4000 ng/mL (Guillette et al., 2022). And finally, snakes showed reduced body condition at serum ΣPFAS concentrations of 100s of ng/mL (Lettoof et al., 2023). The above evidence clearly indicates that low-level chronic exposure leads to diminished health for diverse animal species at tissue concentrations far lower than those needed to produce effects in laboratory acute toxicity studies, and three to four orders of magnitude lower than we observed in the Holloman AFB fauna.

### 4.4. Effects on reproduction

Reproductive and developmental toxicity of several PFAS compounds suggests that reproductive functions for populations at Holloman AFB could be impaired for both birds and mammals. In birds, mother to egg transfer of PFAS occurs predictably, and at the highest rate for PFCAs with longer carbon chains (Jouanneau et al., 2022). In mammals, female mouse livers approximated Holloman AFB levels of PFOS, ∼50k ng/g ww, under an experimental dosage regime that perturbed patterns of gene expression in the placenta, altered corticosterone levels, and reduced body weights (Wan et al., 2020). Humans, mice, and rats have all been shown to have reduced birth weights and poor birth outcomes after developmental exposure to any of a suite of PFAS at tissue concentrations in the single digits ng/g ww (Blake and Fenton, 2020). The breeding bird community at Holloman is undoubtedly subjected to high exposure at early life stages via maternal transfer to eggs (Jouanneau et al., 2022). We tallied 43 bird species for which there is evidence of breeding at or in the vicinity of Holloman Lake, and many additional species likely breed there on an occasional basis. The breeding species tend to rely on resources of the lake, even if nests are not close by. For example, peregrine falcon (*Falco peregrinus*) makes foraging sorties to the lake at all seasons, including from nesting sites that are likely in nearby mountains, ≥ 20 km away. Future studies of the Holloman AFB point source should test for potentially detrimental effects of PFAS on reproduction and development.

### 4.5. Effects on rare species

Several species of conservation concern visit Holloman Lake regularly or occasionally. The Western Snowy Plover (*Charadrius alexandrinus nivosus*) of the interior U.S. is considered “Greatest Conservation Concern” by the U.S. Shorebird Conservation Partnership (shorebirdplan.org) and is a breeding resident around the margins of Holloman Lake, where we observed at least four breeding pairs in 2022. A related species that forages in the same habitat, the Killdeer (*Charadrius vociferus*), proved to be highly contaminated with PFAS (Table 4, Fig.1), as expected for species that forage on the sediment at lake margins (Larson et al., 2018). Several raptor species of conservation concern occur regularly at Holloman Lake: peregrine falcon, prairie falcon (*Falco mexicanus*), bald eagle (*Haliaeetus leucocephalus*), golden eagle (*Aquila chrysaetos*), northern harrier (*Circus hudsonius*), and ferruginous hawk (*Buteo regalis*). These species, other raptors, and locally occurring mammalian carnivore species are expected to experience PFAS trophic magnification, leading to harmful tissue concentrations (see Trophic magnification, below).

### 4.6. Human exposure via hunting, recreation, and livestock

Hunting is a popular activity at Holloman Lake and environs. Oryx, mule deer, pronghorn, javelina, wild boar, jackrabbits, and cottontails are all potentially hunted mammal species occurring in the vicinity. Each of these species should be considered susceptible to PFAS ingestion because they tend to drink free-standing lake water or pumped ground water at stock tanks, and most readily forage near aquatic habitats. Harvest reports for the adjacent White Sands Missile Range Game Management Unit reported 826 oryx hunted during the 2021–2022 season (NMDGF, 2022). Oryx were present in upland scrub areas around Holloman Lake during our sampling efforts and are frequently reported from this locality on iNaturalist.org. Ear-tagged cattle were also present, frequently foraging in the wetlands at the north inflow. PFAS screening of oryx and cattle tissues from this area should be considered a high priority.

In addition to cattle and the seven hunted mammal species, there are at least 41 game bird species that have been recorded at Holloman Lake (Table S2). Non-aquatic, hunted bird species include doves and quail, but even these forage and reproduce in the immediate vicinity of the lake. Hunting of both aquatic and non-aquatic species frequently occurs around the edges of Holloman Lake; duck hunters were present on six of nine days that we visited the lake during the waterfowl hunting seasons of 2021–2022 and 2022–2023. Hunters invariably consume the meat of their quarry after harvest, as required by New Mexico hunting regulations.

Human exposure during outdoor recreational activities is plausible via ingestion of water, soil, or airborne particulates and should be studied further. The sample exhibiting the highest PFOS level encountered in this study (97,000 ng/g) was collected in 1994 at Lagoon G, a wastewater discharge area adjacent to the old golf course, an area utilized heavily for recreational activities by Holloman personnel and families. The samples with the second and third highest contamination levels in 2021–2023 were collected along the Lagoon G outfall canal which drains directly into Holloman Lake. Dispersed camping, recreation, bird watching, and hunting have occurred regularly at Holloman Lake over this nearly three-decade period.

### 4.7. Implications for human health

Routine killing and eating of contaminated game meat provides a pathway for PFAS contamination of human tissues, with damaging effects on health. In pregnant women, PFHxS and PFOA were recently linked to poor birth outcomes, even at very low serum concentrations: geometric means (and standard deviations) of 1.09 (2.30) ng/mL PFHxS and 0.57 (2.31) PFOA, respectively (Taibl et al., 2023).

The European Food Safety Authority (EFSA) has recommended tolerable weekly intake (TWI) levels for contaminated foods; according to these recommendations, the sum of PFOA, PFNA, PFHxS, and PFOS should be no higher than 4.4 ng per kg of body weight per week (EFSA CONTAM Panel et al., 2020). For an adult human weighing 70 kg, this would correspond to a TWI of ∼308 ng; whereas, for a child or adolescent weighing 35 kg, the TWI would be ∼154 ng. We calculated the total amount of meat (pectoral muscle or liver) that could be consumed from Holloman Lake to not exceed these TWIs, based on the mean or maximum values, respectively, for the sum of these four PFAS measured from ducks and coots harvested during 2021–2023. We excluded the common merganser from our calculations because most duck hunters avoid this fishy-tasting species. For pectoral muscle tissue (mean = 3,457 ng/g ww, maximum = 14,150 ng/g ww), the TWI implies a maximum serving size of 89 mg (at mean observed concentration) or 22 mg (at maximum observed concentration) for an adult, or 45 mg (mean) or 11 mg (maximum) for a child. For liver tissue (mean = 14,756 ng/g, maximum = 42,070 ng/g), the TWI would be 21 mg (mean) or 7 mg (maximum) for an adult, or 10 mg (mean) or 4 mg (maximum) for a child. Under the more relaxed standards of the US EPA’s 2016 recommendations, in which the reference level for human consumption of PFOS was 25 ng / kg bw / day (Nolen et al., 2022), a 70-kg adult should consume no more than 12,250 ng PFOS / week which would correspond to 3.54 g duck muscle meat / week. These findings suggest that it would never be safe to eat more than one gram of game bird meat per day from Holloman AFB. For perspective, a U.S. penny weighs 2.5 g. In southeast Australian waterfowl, consumption advisories were issued for duck meat that contained concentrations of PFOS more than two orders of magnitude lower than that which we found at Holloman AFB (Environmental Protection Authority Victoria, 2019; Sharp et al., 2021).

### 4.8. Causes of variation across species

We observed substantial inter-species variation in tissue PFAS profiles (Figs. 1, 3); such variation is expected due to several factors including variable rates of ingestion and elimination. Depending on their ecology and diet, species vary in their relative ingestion of water versus solids and this is expected to alter PFAS carbon-chain length profiles because solid-phase versus aqueous phase substrates differ in degrees of *in situ* PFOS and PFHxS formation from precursors (Anderson et al. 2016). Larson et al. suggested that sediment, not water, is the likely main pathway for avian contamination at federal Air Force Bases contaminated with AFFF; accordingly, a model based on a shoreline foraging charadriiform species, the spotted sandpiper (*Actitis macularia*), predicted the highest levels of tissue contamination (Larson et al., 2018). Consistent with this, we found especially heavy contamination in a juvenile killdeer (MSB:Bird:51309) that we sampled from the shore of Holloman Lake (liver PFOS 23,000 ng/g ww). Even in a shared environment with similar levels of exposure, species vary in tissue concentrations for a range of reasons, including evolved differences in biokinematics that result in different elimination rates (Guruge et al., 2008). Degree of toxicity will depend heavily on these species-specific kinetics (Gomis et al., 2018; Pizzurro et al., 2019). Differences in metabolic rate, liver function, sex, diet, foraging behavior, migration, and other aspects of life history also contribute to variable durations of exposure and variable rates variation in ingestion and elimination (Galatius et al., 2013). The tendency for short-chain PFAAs to accumulate in blue tit eggs but long-chain ones to accumulate in great tit eggs at the same contaminated sites in Belgium highlights the role of species-specific biokinetics (Lasters et al., 2021). Disentangling these drivers of PFAS variation is a challenge for future work, and one that will require substantially more surveillance of wild populations across a range of trophic levels, life history characteristics, and phylogenetic affinities.

### 4.9. Isomer ratios

Fractions of branched versus linear isomers of these three most abundant PFAS compounds revealed interesting patterns with respect to expectations based on manufacturing fractions, reflecting taxon-specific biokinetics, and transfer pathways in the food web that enrich specific isomers (Schulz et al., 2020). Isomeric composition in tissues could have consequences for health effects, as branched isomers of PFOS have been linked with worse pregnancy outcomes than linear isomers (Li et al., 2017). Unexplained enrichment of branched isomers of PFOS in humans suggests that pathways of exposure to PFOS or its precursors and/or pharmacokinetic processes of biotransformation and elimination are not completely understood. Pharmacokinetic processes are further likely to vary among species (Lasters et al., 2021), creating substantial complexity. We found evidence for enrichment in linear isomers of PFOA relative to known typical manufacturing standards in bird, mammal, and plant tissues (Fig. S3, A). Proportions of PFOS isomers fell near expected values based on manufacturing standards for bird muscle and liver and mammal liver; however, mammalian blood was highly enriched in the linear isomer while plant tissues were substantially depleted (Fig. S3, E), emphasizing the importance of taxon-specific exposure and variable pathways of biotransformation and elimination.

### 4.10. PFAS transport in animal tissues

Migrating and wintering waterbirds routinely fly between Holloman Lake and other suitable habitats in the region, providing potential for PFAS-contaminated individuals to be hunted and consumed from uncontaminated sites. The same is generally true for wide ranging mammalian game species. We did not detect individuals heavily contaminated with PFOS, PFOA, or PFHxS among the 10 gamebirds that were screened from uncontaminated control sites. However, one northern pintail from the Middle Rio Grande Valley, central New Mexico, ∼150 km NNW of Holloman Lake, contained among the highest levels that we recorded (850 ng/g ww) of PFNA, a highly toxic, long carbon-chain PFCA (Tables 2, 3, 4). The PFAS profile of this bird (Fig. 3, C) was unique among our samples, suggesting that it had been exposed to PFAS at a site other than Holloman AFB. Clearly more PFAS screening will be needed from broadly dispersed localities, including both contaminated and non-contaminated sites, to test the extent to which PFAS pose a health risk to human hunters and consumers of gamebird meat that was hunted away from point sources. Some species, such as Sandhill Crane (*Antigone canadensis*), regularly stop at Holloman Lake during non-breeding season movements, but are not hunted there; however, they are hunted in neighboring counties, highlighting the possibility that game meat elsewhere could be carrying PFAS contamination that may have accumulated during stopover periods.

### 4.11. Pathways of PFAS movement into aquatic animals

The highest concentrations of PFAS are generally found in the soil and sediment samples from contaminated sites, with levels of PFAS in surface water samples averaging much lower (de Silva et al., 2020). Soil mineral contents, soil PFAS concentrations, and PFAS chain length, strongly influence the level of PFAS uptake by aquatic plants (Pi et al., 2017; Zhang et al., 2020). Zhang *et al*. experimentally demonstrated that long-chain PFAS were integrated into root tissues, while short-chain PFAS were translocated to shoot tissues (Zhang et al., 2020). Trophic magnification of PFAS can occur across all trophic levels but is dependent on characteristics of the PFAS and organismal physiology (Kelly et al., 2009). At the level of primary consumer, benthic macroinvertebrates may accumulate PFAS from herbivory of aquatic plants or directly from exposure to sediment (Brase et al., 2022). For example, in the St. Lawrence River watershed which is exposed to substantial industrial effluent, PFOS in aquatic invertebrates reached as high as 61 ng/g ww, among at least 40 different PFAS detected (Munoz et al., 2022); terrestrial invertebrates near a fluorochemical plant in Belgium measured 28–9000 ng/g ww (D’Hollander et al., 2014). Predatory macroinvertebrates generally accumulate higher PFAS in tissues relative to herbivores, though lifespan and physiological differences may also affect accumulation (Brase et al., 2022). Aquatic animals like ducks accumulate PFAS after feeding on aquatic invertebrates and plants, and by incidental ingestion of sediment and soil. An American wigeon sampled from Holloman Lake had the highest levels of PFAS among any bird in our dataset. This species is primarily herbivorous, feeding mainly on aquatic and terrestrial plants, suggesting that PFAS exposure in gamebirds at Holloman AFB occurs at the lowest trophic levels. Stomach contents of several game bird species in our sampling showed that they were eating large quantities of omnivorous aquatic insects (Hemiptera: Corixidae), a likely pathway for trophic transfer of PFAS (see Arctos.org specimen data linked from Table 1). For many waterbird species, incidental sediment and soil ingestion is thought to cause significant PFAS exposure that is separate from being embedded in a trophic web, (Larson et al., 2018).

### 4.12. Pathways of PFAS movement into terrestrial animals

Ingestion of PFAS in soil and food are the most likely routes of exposure for most terrestrial animals at Holloman AFB; a subset of species that are highly mobile or occur on the lakes edge could also ingest PFAS with surface water. The small mammal community is composed of a suite of species occupying the breadth of environments around Holloman Lake. Species screened for PFAS included: desert granivores such as Merriam’s kangaroo rat (*Dipodomys* merriami), Chihuahuan pocket mouse (*Chaetodipus eremicus*), and western harvest mouse (*Reithrodontomys megalotis*); a generalist, the white-footed mouse (*Peromyscus leucopus*); an omnivore, the Chihuahuan grasshopper mouse (*Onychomys arenicola*); a relatively mesic adapted herbivore, the hispid cotton rat (*Sigmodon hispidus*); and an invasive omnivore, the house mouse. However, there is broad dietary overlap among the arid-adapted Chihuahuan desert rodent species for which seeds and arthropods comprise most of the diet, in proportions that vary seasonally (Hope and Parmenter, 2007). Soil ingestion while foraging or grooming, or dust inhalation likely overshadows exposure through ingestion of water at Holloman Lake for most species. Several of the desert-adapted small mammals in our dataset may not directly ingest water, but rather obtain water from vegetation and invertebrates that they eat. However, a single white-footed mouse (MSB:Mamm:92667) and several house mice (MSB:Mamm:340078, 340121), both species with relatively low tolerance for dehydration (Haines and Schmidt-Nielsen, 1967; MacMillen, 1983) had the highest levels of PFAS among all sampled species (Fig. 1, Table 4). The white-footed mouse was collected from Lagoon G in 1994 (prior to construction of a wastewater treatment plant in 1996). Approximately 1.2 million gallons of domestic and industrial wastewater were discharged to the sewage lagoon area daily, with overflow draining through an outflow canal into Holloman Lake (Amec Foster Wheeler Programs, Inc., 2018). Resampling at this exact location (Lagoon G) was not possible due to denial of access to the military restricted area; however, we were able to collect samples about 1.8 km from Lagoon G in the area of the outfall canal into Holloman Lake. The house mice exhibiting the highest PFAS levels in our contemporary sampling were collected from this location.

It is possible that some of these animals obtained high PFAS levels directly from water consumed from Holloman Lake; however, patterns of PFAS accumulation in water relative to soil suggest that water as a route of exposure may be secondary to dietary and incidental soil exposure (De Silva et al., 2021; Larson et al., 2018). Upland desert rodent species that would not be expected to contact or ingest lake water directly exhibited similar PFAS profiles to rodents of the littoral zone and aquatic birds, with abundant PFOS and PFHxS. By contrast, the four-wing saltbush was high in 6:2 FTS, though ΣPFAS was four orders of magnitude lower than in bird or mammal tissues (Table 2). Soil PFAS are taken up by roots (Stahl et al., 2009), and short-chain PFAS differentially accumulate in plant tissues and invertebrates that feed on them (Ghisi et al., 2019; Groffen et al., 2022). If herbivory were the primary pathway for PFAS ingestion by upland desert rodents, we would expect their PFAS profile to show similarities with that of the four-winged saltbush. We did find a notable excess of 6:2 FTS in rodents relative to aquatic birds, a likely signal of some PFAS transfer via herbivory. Based on the abundance of PFOS and PFHxS as well as the overall similarity of their PFAS profiles to those of aquatic avian secondary consumers at the site, incidental ingestion of soil and inhalation of dust or aerosolized foam from the shoreline are the likely major pathways of PFAS exposure for upland desert rodents. However, more plant sampling is clearly needed to determine whether PFAS uptake by plants and subsequent herbivory represents a significant pathway for PFAS transport to upland desert rodents, particularly for 6:2 FTS.

### 4.13. Trophic magnification

Biomagnification and bioaccumulation of PFOS tend to be most severe in fish-eating upper-trophic predators, particularly air-breathing species (Ankley et al., 2021; Lau et al., 2007). Trophic magnification factors for various PFAS have been estimated to be between ∼1–20 in aquatic systems, ∼2–6 in fish, ∼0.7–7.2 in terrestrial food webs containing birds, and ∼1.1–2.7 in terrestrial food webs with mammals (Kelly et al., 2009; De Silva et al., 2021; Fremlin et al., 2023). PFAS is enriched in longer food chains and, as a result, herbivores tend to have the lowest tissue concentrations (Guckert et al., Miranda et al. 2021). The mean level of PFOS in liver of polar bears (*Ursus maritimus*), a top predator, was as high as 3,270 ng/g ww, even in east Greenland, presumably distant from any point source (Greaves et al., 2012). The primary route of exposure in piscivorous birds is through trophic biomagnification, by which PFAS in sediments and water are translocated into macroinvertebrates, then fish, then piscivores (Larson et al., 2018). PFAS permeates the tissues; it can occur in the bone and skin of mice (Bogdanska et al., 2011) and the skin of ducks (Senversa, 2018) at concentrations similar to blood. PFAS levels in piscivorous birds are highly variable (Kannan et al., 2001) and affected by movement patterns and local PFAS concentrations where foraging occurs. Samples from the piscivorous common merganser collected at Holloman Lake had relatively low PFAS levels, similar to those of control sites.

Interestingly, common merganser is the only piscivorous species that we sampled, and Holloman Lake seems to contain only one species of fish, the mosquitofish (*Gambusia affinis*), an introduced species with very small body size. The stomach of one merganser specimen contained a sunfish (*Leptomis* sp.), which is not known to occur in the lake; thus, we suspect that common mergansers had been using the lake as a rest site between foraging sites in adjacent watersheds, >100 km away. Mergansers were unlike any of the other species of ducks that we tested, each of which consumes invertebrates and vegetative matter. Carnivorous mammals and breeding raptor species that forage on many of the secondary consumer species that we studied here are likely to be at higher risk due to bioaccumulation and trophic magnification of long carbon-chain PFAS (Jouanneau et al., 2020). Bioaccumulation processes tend to be more complex for terrestrial than aquatic food chains (EFSA Contam 2020). Highlighting the dangers for upper trophic predators at Holloman AFB, nearly all tissues that we screened from potential prey species exceeded a benchmark tissue concentration for PFOS of 33 ng/g that was established to protect upper trophic level wildlife species from secondary poisoning (European Union (EU), 2014).

### 4.14. Change over time

Legacy PFAS that were phased out of manufacturing at the turn of the 21^st^ century are anticipated to diminish in the environment over time, both by chemical decomposition and dispersal (Bianchini et al., 2022; Nolen et al., 2022); this is undoubtedly true for Holloman AFB, but with considerable uncertainty about the time scale. Some wildlife monitoring studies have already documented declines in PFOS over time in animal tissues (D’Hollander et al., 2014; Groffen et al., 2017; Jouanneau et al., 2020; Cara et al., 2022; Groffen et al., 2023), although others have found PFOS increases over time (Park et al., 2021) or have found no change in ΣPFAS despite declines in PFOS (Groffen et al., 2023). Wild boar liver tissue at PFAS hotspots in Europe showed evidence of legacy compounds, such as PFOS and PFOA, occurring at tissue concentrations far higher than replacement PFAS compounds that continue to be manufactured (Rupp et al., 2023).

### 4.15. Museum collections and biorepositories for environmental monitoring

This study demonstrates how museum collections of biodiversity research specimens, or biorepositories, are well suited for measuring PFAS contamination in wild populations over space and time. Museum collections have proven essential for ecotoxicological challenges in the past, as exemplified by the discovery that eggshell thinning was caused by the pesticide, DDT (Hickey and Anderson, 1968). Other PFAS studies have also used collections to assess PFAS in ways that would have been impossible without these resources. For example, a study using herbarium specimens showed that pine needles have tracked changes in airborne PFAS levels since the 1960s; the needles sequestered trace amounts of over 70 different PFAS, including new generation ‘replacements’ such as GenX (Kirkwood et al., 2022). In South Korea, birds from a museum collection revealed high levels of contamination in various diurnal and nocturnal birds of prey (Barghi et al., 2018). In the present study, we screened four rodent samples that had been collected during the mid-1990s for other purposes. One specimen proved to be contaminated with PFAS at exceptionally high levels, establishing a nearly three-decade timeline for persistent exposure to these contaminants. Furthermore, the voucher specimens collected for this study in 2021– 2023 are accompanied by frozen tissue samples, ecto- and endoparasites, and online open data, all of which will provide a temporal baseline of the Holloman Lake faunal community for future studies of environmental change. Biorepositories have broadened the range of questions we can ask about PFAS or diverse other aspects of environmental change; for example, species-specific patterns of PFAS exposure, biokinematics, and tolerance will require taxonomically diverse samples that only archival specimens in biorepositories can provide. Combining these contaminant analyses with genomic and epigenetic studies of the specimens or other aspects of their biology provides a powerful framework for assessing the impact of these perturbations on individuals, biotic communities, and ecosystems.

## 5. Conclusions

The biotic communities of Holloman AFB wetlands and surrounding terrestrial habitats are extraordinarily contaminated with PFAS, especially legacy, long carbon-chain forms. This study demonstrates that bird and mammal species living in this environment accumulate those contaminants in their tissues at levels that far exceed those known to affect animal and human health. Our data also highlight the need for expanded monitoring of this desert oasis site; additional data could provide an unusually tractable opportunity to understand PFAS movement through a diverse food web. Biorepositories could provide critical infrastructure for such an effort because they can facilitate sampling that is temporally deep, spatially broad, and taxonomically diverse (Schindel and Cook, 2018). The health of humans who use the Holloman AFB area for hunting or recreation should also be monitored closely; this group of people could be considered sentinels for the overall health of this contaminated environment (Andrews et al., 2023).

The high-dose, acute exposure scenarios on which toxicologists have depended should not be expected to closely approximate chronic exposure scenarios that occur in natural systems, and numerous recent reports have observed harmful effects even at low tissue concentrations; therefore, animal sampling and associated screening data accompanied by measures of health and condition would be a useful extension of research at this site. The Holloman Lake populations of rodents could be models in this respect because of their high and persistent exposure over a period of decades.

The isolated, desert wetlands at Holloman AFB attract migratory birds in abundance at all times of year. To understand how animals might transport PFAS from the site and potential impacts on animal population health, it should be considered a priority to learn where migratory bird populations that use Holloman AFB wetlands spend the remainder of the year. Most bird populations use Holloman Lake seasonally during their migrations to widely dispersed localities in North, Central, and South America. Some species of bats likely to use this site for feeding and drinking are migratory species that winter in Mexico and summer as far north as southern Canada. To understand potential risks to hunters, we need to study population connectivity of game bird and mammal species throughout their annual cycles.

Another major priority for research should be the species-specific dynamics of PFAS bioaccumulation in tissues within and between seasons; for example, it is not yet known whether tissues of wintering birds are less contaminated upon fall arrival than after feeding at the contaminated site for a period of months. For most wild populations that are exposed to PFAS point sources, screening efforts have been inadequate or non-existent. This is especially true for bats, for which no screening data exists to our knowledge. Far more samples and data are needed to establish baselines, parameterize models, and estimate risks to wildlife, livestock, and human health.

## Acknowledgments

We thank the New Mexico Environment Department for funding this study, which was conducted with the collaboration of Daniel B. Stephens & Associates, Inc. (DBS&A). We thank Antonia Androski, Lexi Baca, Sara Brant, Emi Casares, Trinity Casaus, Philip Chaon, Samuel Goodfellow, Danielle Land, Linda Laver, Diana Macias, Tabitha McFarland, Jonathan Mullins, Robert Nofchissey, Colin Peña, Adrienne Raniszewski, Esteban Restrepo Cortes, Spencer Robison, Nella Sanchez Cook, Wildlife Rescue, Inc. of New Mexico, and several anonymous waterfowl hunters for contributions to the sampling, fieldwork, information gathering, laboratory analyses, and specimen preparation. Emily DeArmon identified the fish in the merganser stomach. Natalie Runyon collected the 1994 Holloman specimens. Pat Longmire (NMED) and Amy Ewing (DBS&A) were instrumental in obtaining and managing funds.

## CRediT authorship contribution statement

**Christopher C. Witt:** Conceptualization, Data curation, Funding acquisition, Investigation, Methodology, Project administration, Resources, Supervision, Writing - original draft; **Chauncey R. Gadek:** Data curation, Formal analysis, Methodology, Visualization, Writing - original draft; **Jean-Luc E. Cartron:** Conceptualization, Funding acquisition, Investigation, Methodology, Project administration, Resources, Supervision, Writing - review & editing; **Michael J. Andersen:** Funding acquisition, Supervision, Writing - review & editing; **Mariel L. Campbell:** Investigation, Supervision, Writing - review & editing; **Marialejandra Castro-Farías:** Investigation, Writing - review & editing; **Ethan F. Gyllenhaal:** Investigation, Writing - review & editing; **Andrew B. Johnson:** Conceptualization, Funding acquisition, Investigation, Methodology, Project administration, Supervision, Writing - review & editing; **Jason L. Malaney:** Investigation, Writing - review & editing; **Kyana N. Montoya:** Investigation, Writing - review & editing; **Andrew Patterson:** Data curation, Formal analysis, Investigation, Methodology, Writing - original draft; **Nicholas T. Vinciguerra:** Investigation, Writing - review & editing; **Jessie L. Williamson:** Investigation, Visualization, Writing - review & editing; **Joseph A. Cook:** Conceptualization, Funding acquisition, Investigation, Methodology, Project administration, Resources, Supervision, Writing - original draft; **Jonathan L. Dunnum:** Conceptualization, Funding acquisition, Investigation, Methodology, Project administration, Supervision, Writing - original draft.

---

**Table S1.** Proportion of samples that exceeded reporting limits for each combination of taxonomic class, tissue type, and sampling locality, and for each of 17 PFAS and six isomeric constituents that were assayed.

| PFAS | bird |  |  |  | mammal |  |  |  | plant |
| --- | --- | --- | --- | --- | --- | --- | --- | --- | --- |
|  | liver |  | muscle |  | blood |  | liver |  | leaf/stem |
|  | control | Holloman | control | Holloman | control | Holloman | control | Holloman | Holloman |
|  | n = 1 | n = 24 | n = 9 | n = 15 | n = 1 | n = 9 | n = 4 | n = 34 | n = 2 |
| Perfluorobutanoic acid (PFBA) | 0.00 | 0.46 | 0.00 | 0.40 | 0.00 | 0.67 | 0.25 | 0.15 | 1.00 |
| Perfluoropentanoic acid (PFPeA) | 0.00 | 0.00 | 0.00 | 0.00 | 0.00 | 0.33 | 0.00 | 0.09 | 1.00 |
| Perfluorohexanoic acid (PFHxA) | 0.00 | 0.00 | 0.00 | 0.00 | 0.00 | 0.11 | 0.00 | 0.12 | 1.00 |
| Perfluoroheptanoic acid (PFHpA) | 0.00 | 0.17 | 0.00 | 0.07 | 0.00 | 0.44 | 0.00 | 0.15 | 0.50 |
| L-Perfluorooctanoic acid | 1.00 | 0.71 | 0.00 | 0.87 | 0.00 | 0.33 | 0.50 | 0.38 | 0.00 |
| Br-Perfluorooctanoic acid | 0.00 | 0.21 | 0.00 | 0.07 | 0.00 | 0.22 | 0.25 | 0.18 | 0.00 |
| Total PFOA | 1.00 | 0.71 | 0.00 | 0.87 | 0.00 | 0.33 | 0.50 | 0.38 | 0.00 |
| Perfluorononanoic acid (PFNA) | 1.00 | 0.88 | 0.11 | 0.93 | 0.00 | 1.00 | 0.50 | 0.74 | 0.00 |
| Perfluorodecanoic acid (PFDA) | 1.00 | 0.92 | 0.00 | 0.93 | 0.00 | 0.89 | 0.00 | 0.65 | 0.00 |
| Perfluoroundecanoic acid (PFUnA) | 1.00 | 0.71 | 0.00 | 0.13 | 0.00 | 1.00 | 0.00 | 0.53 | 0.00 |
| Perfluorobutanesulfonic acid (PFBS) | 0.00 | 0.04 | 0.00 | 0.00 | 0.00 | 0.11 | 0.00 | 0.03 | 1.00 |
| Perfluoropentanesulfonic acid (PFPeS) | 0.00 | 0.54 | 0.00 | 0.80 | 0.00 | 0.67 | 0.00 | 0.15 | 0.00 |
| Br-Perfluorohexanesulfonic acid | 0.00 | 0.46 | 0.00 | 0.73 | — | — | 0.25 | 0.53 | 0.50 |
| L-Perfluorohexanesulfonic acid | 0.00 | 0.88 | 0.00 | 0.87 | — | — | 0.50 | 0.82 | 1.00 |
| Total PFHxS | 0.00 | 0.88 | 0.00 | 0.87 | 0.00 | 1.00 | 0.50 | 0.82 | 1.00 |
| Perfluoroheptanesulfonic acid (PFHpS) | 0.00 | 0.71 | 0.00 | 0.87 | 0.00 | 1.00 | 0.50 | 0.74 | 0.00 |
| L-Perfluorooctanesulfonic acid | 1.00 | 0.92 | 0.33 | 1.00 | 1.00 | 1.00 | 0.75 | 0.97 | 1.00 |
| Br-Perfluorooctanesulfonic acid | 0.00 | 0.83 | 0.00 | 0.93 | 0.00 | 1.00 | 0.75 | 0.91 | 1.00 |
| Total PFOS | 1.00 | 0.96 | 0.33 | 1.00 | 1.00 | 1.00 | 0.75 | 0.97 | 1.00 |
| 4:2 FTS | 0.00 | 0.00 | 0.00 | 0.00 | 0.00 | 0.00 | 0.00 | 0.00 | 0.00 |
| 6:2 FTS | 0.00 | 0.13 | 0.00 | 0.00 | 0.00 | 0.22 | 0.50 | 0.35 | 1.00 |
| 8:2 FTS | 0.00 | 0.79 | 0.00 | 0.80 | 0.00 | 0.33 | 0.25 | 0.44 | 0.00 |
| 10:2 FTS | 0.00 | 0.04 | 0.00 | 0.00 | 0.00 | 0.00 | 0.00 | 0.06 | 0.00 |

**Table S2.** Species of birds and mammals that are hunted for their meat and are known to occur at Holloman Lake. Bird occurrence data and taxonomy follow ebird.org. Mammal taxonomy follows the Mammal Diversity Database, mammaldiversity.org (Burgin et al., 2018). These are species that are regularly and legally hunted and eaten by people, providing pathways for PFAS entry to human tissues.

| Class | Family | Common Name | Scientific Name |
| --- | --- | --- | --- |
| Bird | Anatidae | Fulvous Whistling-Duck | *Dendrocygna bicolor* |
| Bird | Anatidae | Snow Goose | *Anser caerulescens* |
| Bird | Anatidae | Canada Goose | *Branta canadensis* |
| Bird | Anatidae | Wood Duck | *Aix sponsa* |
| Bird | Anatidae | Blue-winged Teal | *Spatula discors* |
| Bird | Anatidae | Cinnamon Teal | *Spatula cyanoptera* |
| Bird | Anatidae | Northern Shoveler | *Spatula clypeata* |
| Bird | Anatidae | Gadwall | *Mareca strepera* |
| Bird | Anatidae | Eurasian Wigeon | *Mareca penelope* |
| Bird | Anatidae | American Wigeon | *Mareca americana* |
| Bird | Anatidae | Mallard | *Anas platyrhynchos* |
| Bird | Anatidae | Mexican Duck | *Anas diazi* |
| Bird | Anatidae | Northern Pintail | *Anas acuta* |
| Bird | Anatidae | Green-winged Teal | *Anas crecca* |
| Bird | Anatidae | Canvasback | *Aythya valisineria* |
| Bird | Anatidae | Redhead | *Aythya americana* |
| Bird | Anatidae | Ring-necked Duck | *Aythya collaris* |
| Bird | Anatidae | Greater Scaup | *Aythya marila* |
| Bird | Anatidae | Lesser Scaup | *Aythya affinis* |

| **Class** | **Family** | **Common Name** | **Scientific Name** |
| --- | --- | --- | --- |
| Bird | Anatidae | Surf Scoter | *Melanitta perspicillata* |
| Bird | Anatidae | White-winged Scoter | *Melanitta deglandi* |
| Bird | Anatidae | Black Scoter | *Melanitta americana* |
| Bird | Anatidae | Long-tailed Duck | *Clangula hyemalis* |
| Bird | Anatidae | Bufflehead | *Bucephala albeola* |
| Bird | Anatidae | Common Goldeneye | *Bucephala clangula* |
| Bird | Anatidae | Hooded Merganser | *Lophodytes cucullatus* |
| Bird | Anatidae | Common Merganser | *Mergus merganser* |
| Bird | Anatidae | Red-breasted Merganser | *Mergus serrator* |
| Bird | Anatidae | Ruddy Duck | *Oxyura jamaicensis* |
| Bird | Odontophoridae | Scaled Quail | *Callipepla squamata* |
| Bird | Odontophoridae | Gambel's Quail | *Callipepla gambelii* |
| Bird | Columbidae | Rock Pigeon | *Columba livia* |
| Bird | Columbidae | Eurasian Collared-Dove | *Streptopelia decaocto* |
| Bird | Columbidae | White-winged Dove | *Zenaida asiatica* |
| Bird | Columbidae | Mourning Dove | *Zenaida macroura* |
| Bird | Rallidae | Virginia Rail | *Rallus limicola* |
| Bird | Rallidae | Sora | *Porzana carolina* |
| Bird | Rallidae | American Coot | *Fulica americana* |
| Bird | Gruidae | Sandhill Crane | *Antigone canadensis* |
| Bird | Scolopacidae | Wilson's Snipe | *Gallinago delicata* |
| Mammal | Bovidae | Oryx | *Oryx gazella* |
| Mammal | Cervidae | Mule Deer | *Odocoileus hemionus* |
| Mammalia | Antilocapridae | Pronghorn | *Antilocapra americana* |
| Mammal | Tayassuidae | Javelina | *Dicotyles tajacu* |
| Mammal | Suidae | Wild Boar | *Sus scrofa* |
| Mammal | Leporidae | Desert Cottontail | *Sylvilagus audubonii* |
| Mammalia | Leporidae | Black-tailed Jackrabbit | *Lepus californicus* |

**Fig. S1.**
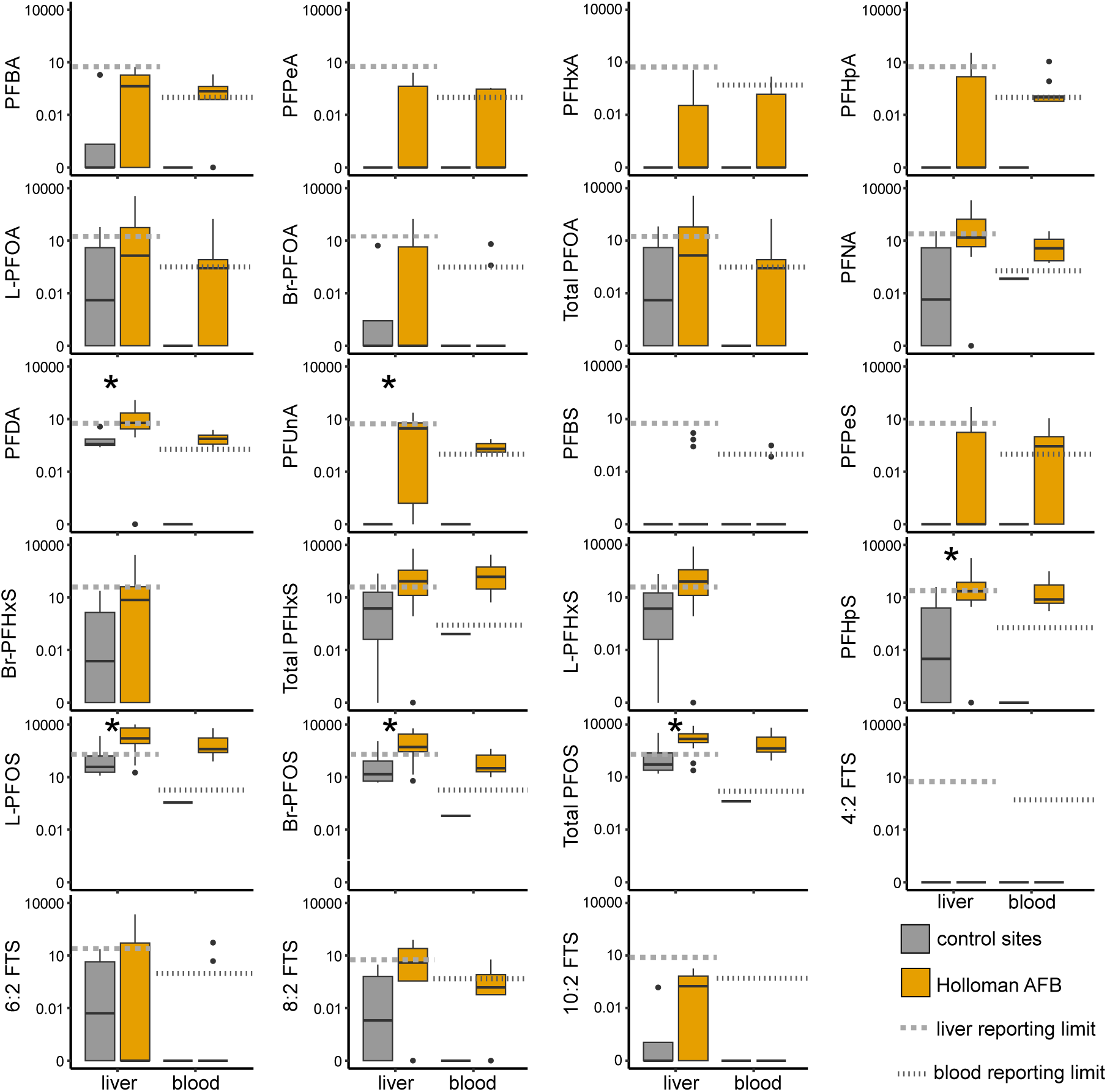
Comparison of PFAS concentrations between mammal tissue samples from Holloman Lake (orange) and control sites (gray), for blood (ng/mL) and liver (ng/g), respectively. Gray dotted lines and dashed lines indicate mean reporting limit thresholds per compound for blood and liver, respectively. Asterisks indicate significant differences between concentrations calculated from Kruskal-Wallis tests for liver samples only (*p< 0.05). Because we had only a single blood sample from a control site, we did not run significance tests for blood; however, the control-site blood sample did not exceed the reporting limit for any compound, in strong contrast to the Holloman AFB blood samples.

**Fig. S2.**
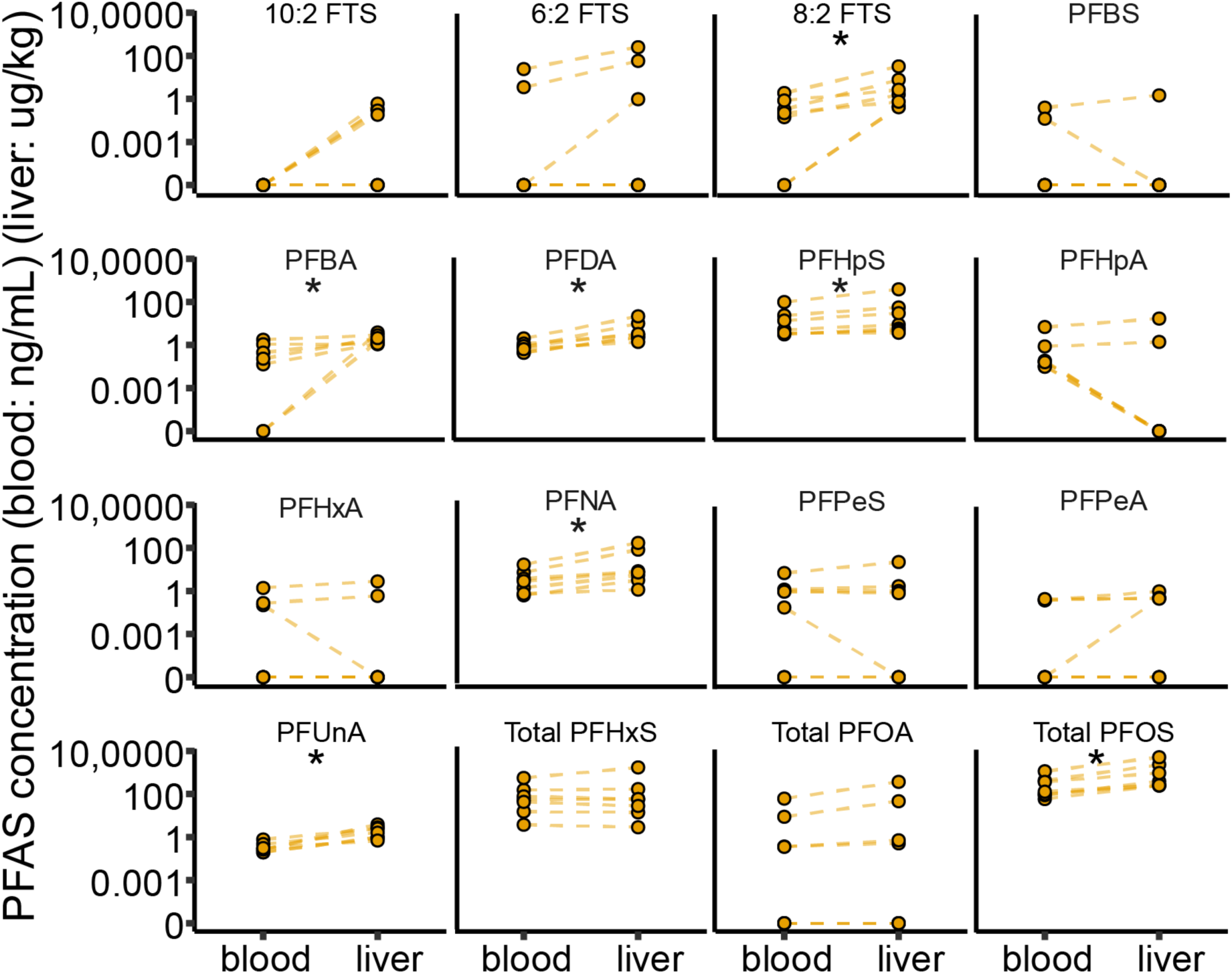
Between tissue comparison for blood versus liver of small mammals. Points are blood sample values measured in ng/mL and liver values in ng/g units. Lines connect points from the same animal. Steeper lines connecting tissues within individuals represent larger between tissue differences in PFAS concentrations. Asterisks indicate significant differences in PFAS concentrations between tissues based on Wilcoxon Signed Rank tests using normal approximation (*p < 0.05).

**Fig. S3.**
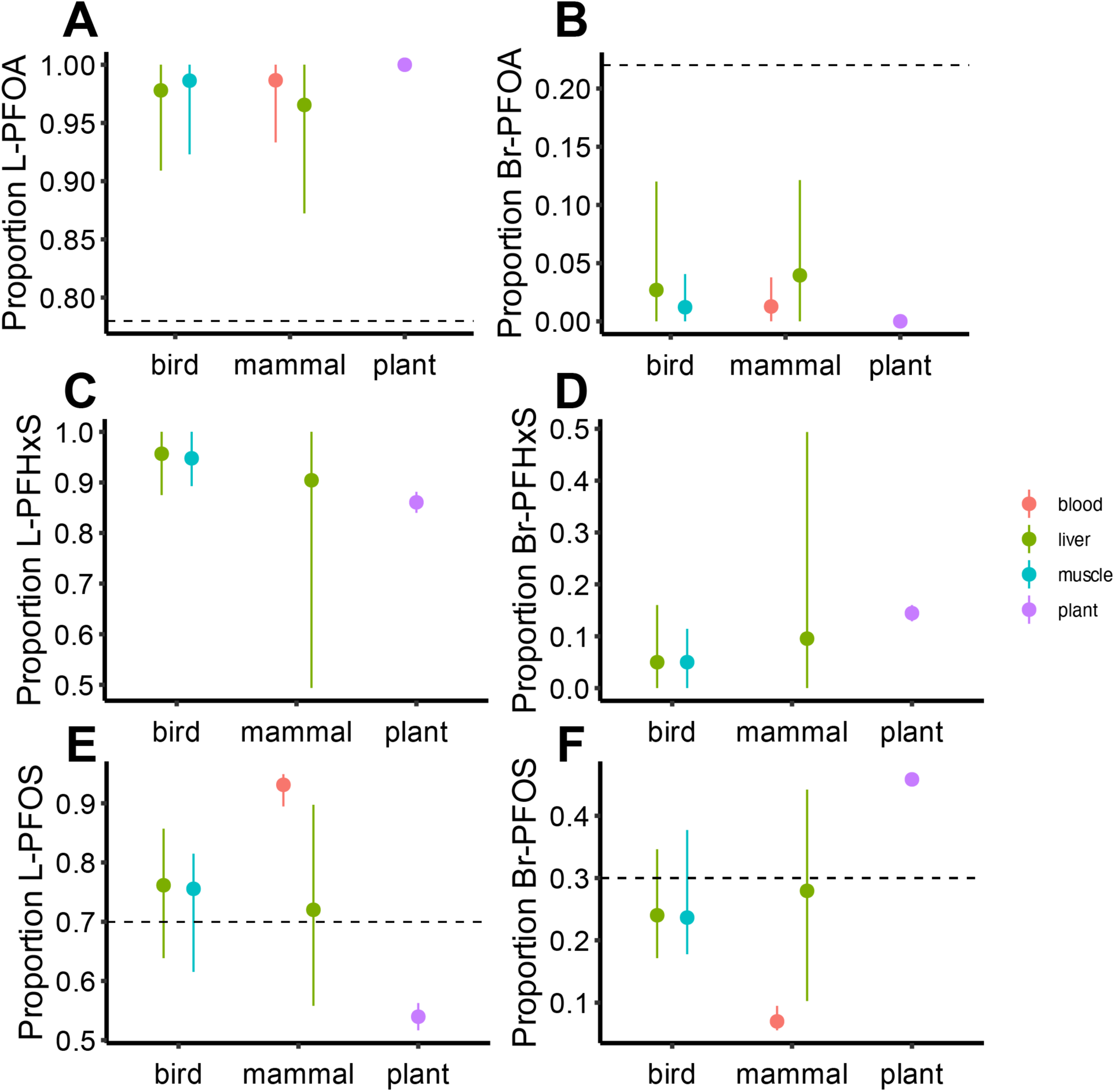
Proportions (points) and range (vertical lines) of linear (L-) and branched (Br-) isomers of PFOA, PFHxS, and PFOS for bird (muscle and liver), mammal (blood and liver) and plant (stems and leaves). Dashed horizontal lines indicate typical manufacturing isomeric proportions. (A, C, E) Linear proportions. (B, D, F) Branched proportions.

